# HuBIE: The Human Blood Immunome Encyclopedia Of TCRs and BCRs in Bloodstream Infections and Cancer

**DOI:** 10.1101/2025.10.22.678684

**Authors:** Vahid Akbari, Alexandra Morgan, Michie Yasuda, Ulrich Schlecht, Sylvie McNamara, Hossein Asgharian, Christopher Tam, Rena Adachi, Elisa Contreras, Zhipei Gracie Du, Sandra Siemann, Hahn Zhao, Jeyashree Ashok Balasubramanian, Diane Balallo, Devanshee Sanghvi, Tarini Shankar, Sanjucta Dutta, Stefan Riedel, Stefanie Mattson, Daniel Burukhin, Florian Rubelt, Sowmi Utiramerur, Dinesh Kumar, Hamid Mirebrahim, Ramy Arnaout

**Author notes:** Equal Contribution (Vahid Akbari, Alexandra Morgan, Michie Yasuda). To whom correspondence should be addressed at or.

## Abstract

T and B cells are central to adaptive immunity, where they identify and neutralize foreign antigens and cancer neo–antigens. Large–scale elucidation of T– and B–cell receptors (TCRs and BCRs) through immune–repertoire sequencing promises novel diagnostics, prognostic markers, and therapeutic strategies. However, progress is hampered by small cohort sizes, a lack of real–world patient diversity, and heterogeneous sample processing, impeding cross–study comparability. To overcome these limitations, here we introduce the Human Blood Immunome Encyclopedia (HuBIE), comprising immune–repertoire data from 2,614 samples collected from 1,941 participants. The cohort includes a range of bloodstream infections, several cancer types, and control participants, with many individuals providing longitudinal samples. We employed Roche’s immune receptor Primer Extension Target Enrichment (immunoPETE) platform to perform simultaneous targeted sequencing of T–cell receptor β chains (TRB), T–cell receptor δ chains (TRD), and immunoglobulin heavy chains (IGH), thereby profiling TCRs and BCRs in all participants. We provide a comprehensive description of immune–repertoire diversity in cancer and bloodstream infections and examine variations across demographic variables such as age and race. We find significant differences in TRB and IGH composition across ethnic groups, and show that the fall in repertoire diversity with age follows distinct patterns for TRB, TRD, and IGH and is accompanied by age-related differences in VJ gene usage. Finally we demonstrate that greater immunological diversity is associated with improved survival but only for elderly participants. HuBIE thus constitutes a valuable resource for the immune–repertoire community, enabling large–scale mapping of the human immunome to accelerate development of diagnostics, prognostic biomarkers, and innovative therapeutic strategies.

## Introduction

The human adaptive immune system is built around two highly diverse lymphocyte lineages: T cells and B cells. These cells are uniquely equipped to recognize and react to an enormous variety of antigens presented by pathogens and cancer cells. Much of this versatility stems from two mechanisms: somatic recombination and junctional diversity^1,2^. Somatic recombination creates novel T–cell receptors (TCRs) and B–cell receptors (BCRs) by randomly combining hundreds of variable (V), diversity (D), and joining (J) gene segments. Junctional diversity further expands the repertoire by inserting random nucleotides at the V–D and D–J junctions. Combined, these processes generate on the order of hundreds of billions of distinct receptor sequences in a single individual at any given point in time, often with very little overlap between different people^1–3^. BCRs also undergo class–switch recombination and somatic hypermutation, which further increases sequence diversity^4^. Consequently, the complete set of unique TCR and BCR sequences—referred to as the immune repertoire or immunome—is highly dynamic, evolving in response to disease and other physiological conditions, and thus serves as a valuable biomarker for diagnosis, prognosis, and therapeutic monitoring^4–9^.

Immune–repertoire sequencing has become an invaluable tool for advancing our understanding of immune responses across many fields, particularly in two critical clinical areas: cancer and bacterial bloodstream infections^9–12^. In cancer, the technique enables detailed profiling of TCR and BCR diversity, illuminating tumor–immune interactions that are crucial for the development of personalized immunotherapies and for assessing treatment outcomes. In bacterial bloodstream infections, immune–repertoire sequencing reveals how the host recognizes and combats invasive pathogens, thereby informing vaccine design and antimicrobial–strategy development^5,6,12–16^. By delivering high–resolution data on immune repertoires, this approach fills important knowledge gaps in each domain and paves the way for innovative diagnostic, prognostic, and therapeutic solutions in these challenging clinical settings. Nevertheless, immune repertoire research is frequently limited by reliance on immune–receptor inference from whole–transcriptome data, small cohort sizes, the availability of only BCR or only TCR data, insufficient sample diversity, and heterogeneous processing technologies—all of which can introduce bias and impede cross–study comparability.

We introduce the Human Blood Immunome Encyclopedia (HuBIE), a novel, large–scale collection of T–cell–receptor (TCR) and B–cell–receptor (BCR) repertoires. HuBIE comprises 2,614 blood–derived samples obtained from 1,941 real–world patients who sought care for bloodstream infection, cancer, or both—leading causes of morbidity and mortality worldwide. It is distinguished not only by its scale—hundreds of participants per condition and tens of thousands of unique sequences per individual—but also by its exclusively real–world, all–comer cohort, which provides a broadly generalizable benchmark that captures authentic population heterogeneity, treatment patterns, and outcomes. All samples were processed with Roche’s immune receptor Primer Extension Target Enrichment (immunoPETE) platform, which enables simultaneous sequencing of the CDR3 segment of TCR beta (TRB) and delta (TRD) chains together with immunoglobulin heavy (IGH) chains as well as determination of V and J gene segment identities^5,6,17,18^. Our analysis reproduces several previously reported observations on immune repertoires in both cancer and infectious–disease settings. Through the cohort’s scale and breadth, we also identify age– and race–associated differences in repertoire composition, longitudinal alterations within participants, and features linked to survival. For every participant, HuBIE provides detailed IGH, TRB, and TRD profiles and is accompanied by extensive clinical annotation, including demographics, comorbidities, treatments, and survival status. This combination of deep immunome data and rich clinical metadata fills gaps in existing resources, supporting personalized immunotherapy research and facilitating the development of diagnostics, prognostics, and innovative therapeutic approaches across a broad cross-section of the population. Moreover, the size of this dataset creates unique opportunities to train and validate machine–learning models for a variety of biomedical applications.

## Results

### Overview of the Cohort and Immune-repertoire sequencing

We set out to profile T–cell–receptor (TCR) and B–cell–receptor (BCR) repertoires across a broad spectrum of human disease states. Using Roche’s immunoPETE platform, we simultaneously sequenced recombined TCR and BCR genes for every specimen. The study cohort comprises 1,941 individuals contributing a total of 2,614 blood samples (Fig. 1a; Supplementary Table 1). Longitudinal sampling is available for 419 participants, yielding 1,092 time–point samples (Fig. 1b). Samples were classified into five clinical groups: (i) “infection,” participants with culture–confirmed bloodstream infection (16.5 % of samples); (ii) “cancer,” participants with an electronic–medical–record (EMR)–confirmed cancer diagnosis (50.5 %); (iii) “culture-negative,” participants suspected of bloodstream infection with negative blood cultures (14.2 %); (iv) “cancer + infection,” participants meeting both cancer and infection criteria (7.0%); and (v) “control,” participants without infection or cancer (11.9 %) (see Methods). Comprehensive demographic data—including sex, age, and self–reported race (white, Black, Asian, Hispanic, and Native American)—are recorded for the entire cohort (Fig. 1a). The sex distribution is balanced, with roughly half of the participants male and half female. There were older adults than younger ones, reflecting the underlying demographics of bloodstream infection and cancer. Aside from a limited number of Native American participants, the cohort is racially and demographically diverse, with strong representation across all major groups. This diversity ensures that each age stratum contains substantial representation of both sexes and of all racial categories (Fig. 1a; Supplementary Table 1).

**Figure 1.**
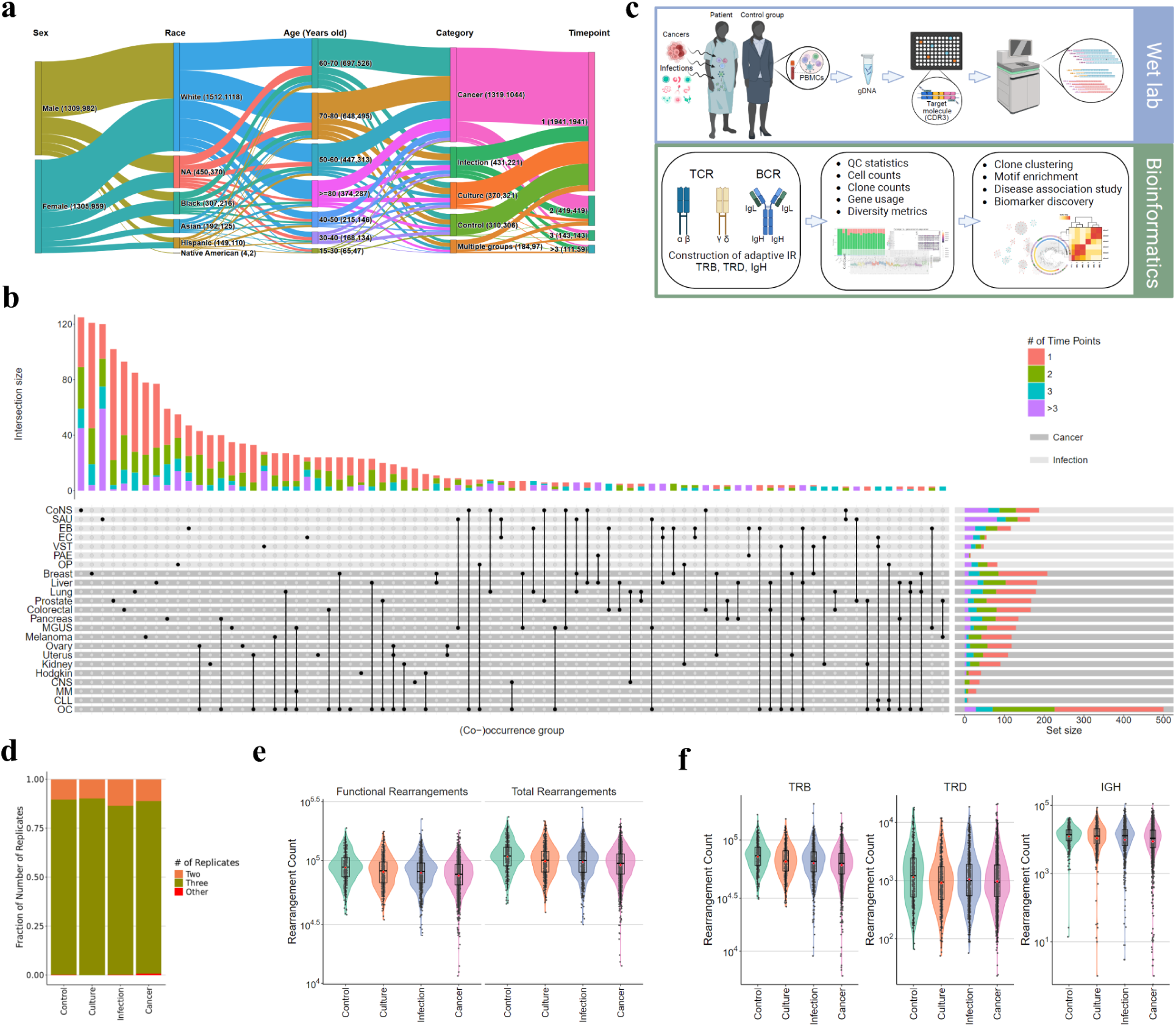
Overview of the cohort composition and categories. **a)** Sankey diagram of (number of samples, number of participants) by demographic, study group, and time point number. Note that the total number of participants in the cohort is 1,941. However, the sum of the number of participants in the age, category, and timepoint columns exceeds 1,941 because longitudinal samples belong to multiple timepoints and the category or age group for some of these participants has also changed at some timepoints. **b)** UpSet plot of the number of samples for each bacteria and cancer type illustrating co-occurrences for all (co-)occurrence groups with ≥3 intersections. The bars are colored to show the proportion of samples based on the number of timepoints including samples with 1, 2, 3, or more than 3 timepoints. About 63% (1,210/1,934) of cancer or infection samples belong to only one cancer or infection subtype (considering other pathogens and other cancers as separate subtypes), 26% belong to multiple cancer subtypes, 1.7% belong to multiple infection subtypes, and 9.5% belong to both cancer and infection subtypes. In cancer subtypes the abbreviations are CNS (Central nervous system), MGUS (Monoclonal gammopathy of undetermined significance), CLL (Chronic lymphocytic leukemia), MM (Multiple myeloma), and OC (Other cancers). In infection subtypes the abbreviations are EB (Enterobacterales), EC (Enterococci), CoNS (Coagulase-negative staphylococci), VST (Viridans streptococci), PAE (Pseudomonas aeruginosa), SAU (Staphylococcus aureus), and OP (Other pathogens). **c)** immunoPETE workflow. PBMC samples are collected from enrolled participants, DNA is extracted, libraries are prepared and sequenced (using the NovaSeq 6000 platform), and sequence data processed using the immunoPETE bioinformatics pipeline to call functional vs. non-functional TCR and BCR clonotypes and assign V and J gene segments. **d)** Stacked bar graph of the number of immunoPETE replicates in each category. **e)** Total number of VDJ rearrangements and number of functional VDJ rearrangements in each sample in each group. **f)** Total number of functional VDJ rearrangements of each chain in the samples of each group.

immunoPETE has previously been used to profile and study the immune repertoire^5,6,17,18^. Applying this platform, we sequenced immune–receptor DNA from blood specimens of HuBIE participants (Fig. 1c). The majority of samples (83.7%) were run in triplicate; 15.8% were processed as duplicates because of lower DNA availability (Fig. 1d). In total we detected ∼261 million VDJ rearrangements, of which >213 million (82%) were functional. A detailed cohort breakdown yielded 34.5 million rearrangements from control, 37.8 million from culture, 44.5 million from infection, 126.6 million from cancer, and 17.6 million from cancer + infection specimens, corresponding to per–sample means ± SD of 111,251 ± 30,345, 102,230 ± 28,493, 103,203 ± 29,014, 95,950 ± 27,323, and 95,554 ± 26,952 rearrangements, respectively (Fig. 1e). Across all groups, ∼82 % of rearrangements were functional (in–frame VDJ recombination without stop codons), a proportion that aligns with the ∼14 % non–productive CDR3 frequency reported by Murugan et al. (Fig. 1e)^19^. Because non–functional rearrangements do not generate productive T– or B–cell receptors, downstream analyses were confined to functional events, which comprised 172.0 million TRB, 4.2 million TRD, and 37.0 million IGH sequences. On a per–sample basis, TRB chains accounted for ∼81 % of functional VDJ events, whereas IGH and TRD chains contributed ∼17 % and ∼2 %, respectively (Fig. 1f), consistent with known peripheral–blood frequencies of these lymphocyte subsets^20,21^. The mean ± SD counts per sample were 65,785 ± 21,203 for TRB, 1,622 ± 1,955 for TRD, and 14,167 ± 11,025 for IGH (Fig. 1f). Collectively, our cohort together with the immunoPETE sequencing data constitute a comprehensive resource that integrates detailed demographics, disease phenotypes, and simultaneous TCR and BCR clonotype information for future immunological research.

### Repertoire Patterns by Sex, Age, and Race

Immunological diversity is biologically important and may have diagnostic or prognostic value^3,15,16,22–24^. We applied Hill’s diversity–number framework to compute the “D-number” form of four standard metrics: species richness (corresponding to *D*^0^), Shannon entropy (*D*^1^), Simpson’s index (*D*^2^), and the Berger–Parker clonality index (*D*^∞^)^23,25^. D numbers express these metrics in the same units: the effective number of clonotypes. The superscript denotes the weight given to clone size: *D*^0^ measures richness regardless of clone size, whereas *D*^∞^ reflects the space left by the single largest clone. Small-superscript metrics (*D*^0^, *D*^1^) are sensitive to the total number of rearrangements or sequencing reads obtained (Supplementary Fig. 1a), a statistical sampling bias that can generate misleading conclusions if not addressed^23^. In our dataset, even state–of–the–art correction methods failed to fully remove this bias (Supplementary Fig. 1a)^23,26^. Therefore we down–sampled every sample to an identical number of per-cell productive rearrangements before calculating diversity (see Methods; Supplementary Fig. 1a).

HuBIE’s cohort spans a wide demographic spectrum representing both sexes and participants 18 to >90 years old (median = 66 years old; mean ± SD = 64 ± 15.4 years old). The self–reported racial composition broadly mirrors the United States population (Fig. 1a). These features provide a unique opportunity to examine how IGH, TRB and TRD repertoire diversity varies with age, race, and sex. Previous work has shown that immunological diversity declines with age^27,28^; this age–related reduction in diversity is also linked to altered disease susceptibility and immune–response outcomes^5,28^. For example, we previously reported a pronounced narrowing of both TRB and TRD diversity in COVID–19 participants older than 50 years, a pattern that was absent in younger individuals^5^. This observation suggests that loss of T–cell diversity may underlie the increased disease severity seen in older participants.

We observed a uniform decrease with increasing age across all diversity metrics (from *D*^0^ to *D*^∞^) for IGH, TRB and TRD in both sexes (Fig. 2a, Supplementary Fig. 2a). This reflects a reduction in the total number of unique clones (*D*^0^) and an expansion of the relative size of the largest clones (*D*^∞^). Importantly, the trajectory of this decline varies markedly among the three receptor chains, as follows.

**Figure 2.**
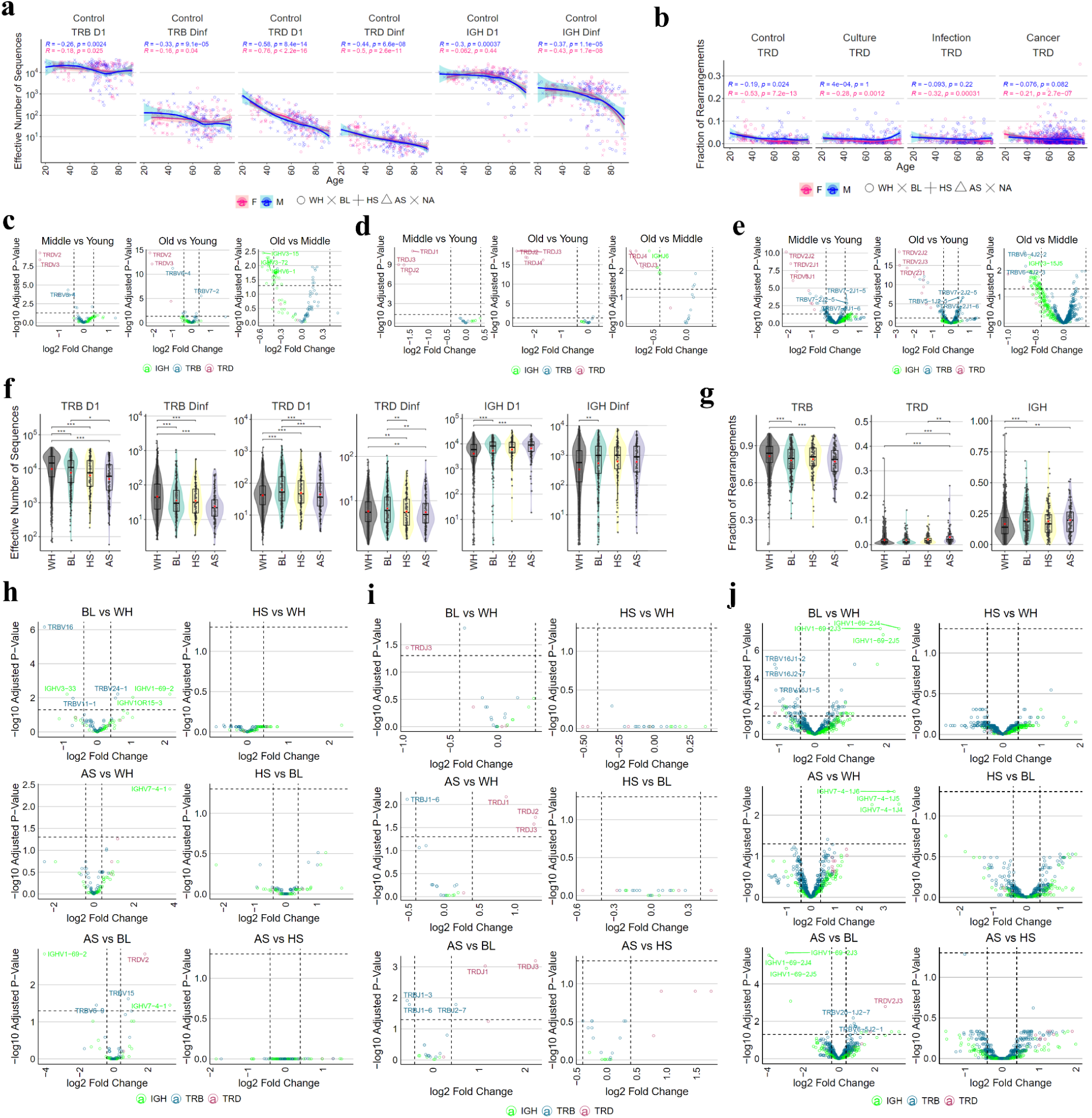
Immune repertoire patterns with demographics. **a)** Spearman correlations between the diversity measures *D*^1^ and *D*^∞^ with age for each chain type in control group. **b)** Spearman correlations between TRD chain fractions with age in different conditions. **c-e)** V (c), J (d), and VJ (e) Gene usage differential analysis between age groups in control samples. **f)** *D*^1^ and *D*^∞^ for each chain type in each race across the cohort. **g)** Fraction of each chain type in the immune repertoire across different races. **h-j)** V (h), J (i), and VJ (j) Gene usage differential analysis between race groups in control samples. WH, white; BL, black; HS, hispanic; AS, asian. In race analysis, native Americans (AM) are excluded as there were very few AM samples. In box and violin plots statistical test was performed using a robust linear regression model (with covariate adjustment) and p values are FDR or Benjamini-Hochberg adjusted. * represents p value<0.05; ** p<0.01; *** p<0.001. Gene usage analysis was performed using limma-voom (with covariate adjustment) and p values are Benjamini-Hochberg adjusted. For the sake of visualization in volcano plots, only top 3 up- or down-used genes are labeled.

For IGH, the trend is driven primarily by growing inter–individual variability with age: older individuals are increasingly likely to exhibit markedly lower diversity than their peers, a pattern that becomes especially evident after age 60 (Fig. 2a, Supplementary Fig. 2a). *D*^0^ and *D*^1^ decline more slowly than *D*^∞^, showing that the total clone count remains relatively stable while clonality (the dominance of the largest clone) rises. These observations are compatible with two dynamical scenarios: (i) IGH diversity stays roughly constant for decades and then collapses suddenly for most participants; or (ii) IGH diversity declines continuously, but the rate is dramatically faster in a subset of participants. Our results refute two alternative hypotheses: (i) a uniform, gradual decline across the entire population, and (ii) the existence of widely divergent baseline diversities at birth.

By contrast, TRB does not show an age–related rise in heterogeneity (Fig. 2a, Supplementary Fig. 2a). Both *D*^0^, *D*^1^, and *D*^∞^ decrease for TRB, but the decline is modest compared with IGH, and inter–individual heterogeneity stays approximately constant across ages. Thus, TRB diversity appears to reflect distinct baseline levels among individuals, followed by a mild age–related decrease. Finally, TRD follows a third trajectory: both *D*^0^ and *D*^∞^ decline steadily with age, mirroring the well–documented reduction in γδ–T–cell production (Fig. 2a, Supplementary Fig. 2a)^27,29^. Consistent with this decline, age correlates negatively with the proportion of TRD sequences and, conversely, shows a weak positive correlation for TRB (Fig. 2b, Supplementary Fig. 2b).

Earlier work documented an age–related decline in the similarity of TRB repertoires between individuals; our substantially larger HuBIE cohort reproduces this pattern and extends it to TRD and IGH (Supplementary Fig. 3a, 4a)^30^. We calculated pairwise repertoire overlap for three age brackets—young (< 40 yr), middle–aged (40–65 yr) and old (> 65 yr)—using a maximum Hamming distance of 1 to define near–identical CDR3 sequences (Supplementary Fig. 3a, 4a). Young donors yielded the highest overlap scores for every chain, with the strongest effect observed for TRD. Differential usage of V, J, and V–J gene pairs was then examined across the same age groups, revealing clear age–dependent shifts in gene utilization (Fig. 2c–e). The most striking alterations occurred in the transition from youth to middle age for TRD: V, J and V–J combinations that dropped markedly in the middle–aged versus young comparison continued to decline further in the old versus young comparison. This pattern indicates that the progressive loss of TRD diversity stems from a skewed V–J representation rather than an even erosion of all TRD sequences. In contrast, TRB and IGH repertoires changed far less dramatically. TRB showed increase in several V–J pairs while decrease in several other V-J pairs in middle age and old compared to young individuals. Many IGH V, J and V–J combinations were used less frequently when participants moved from middle age to old age (Fig. 2c–e). Finally, CDR3 charge and length remained essentially unchanged with age for IGH, TRB and TRD (Supplementary Fig. 5).

The large size and diverse demographic composition of HuBIE allowed for a comparative analysis of immune repertoires across self-described racial groups. We applied a robust linear–regression model that adjusts for age and sex effects to compare the racial groups. This analysis uncovered statistically significant differences between Asians and whites, with whites exhibiting greater TRB diversity and lower IGH diversity; Black and Hispanic participants displayed intermediate values (Fig. 2f; Supplementary Fig. 6a). These differences were evident at both ends of the Hill number spectrum (*D*^1^ and *D*^∞^), indicating that whites harbor larger numbers of smaller TRB clones while possessing fewer but larger IGH clones. Consistent with these findings, whites had the largest fraction of TRB sequences and the lowest IGH fraction, whereas Asians showed the opposite pattern (Fig. 2g). In parallel, Asians exhibited the highest fraction of TRD rearrangements, and whites the lowest (Fig. 2g). These racial trends persisted across multiple clinical conditions but reached statistical significance primarily in the cancer group (Supplementary Fig. 6b-f). We also identified statistically significant differences in V, J, and V–J gene–pair usage, particularly among Asians, whites, and Blacks (Fig. 2h-j). Notably, the IGH V7-4-1 gene paired with any of J4-6 gene was used more frequently in Asians than whites. Conversely, IGH V1-69-2 paired with any of J3-5 occurred less often in Asians and whites compared with Blacks. Finally, modest yet statistically significant variations in CDR3 charge and length were observed across racial groups (Supplementary Fig. 7).

### Exploring Immune Repertoire in Control vs. Disease Groups

We employed a robust linear–regression model that adjusts for age, sex, and self–identified race to compare immune–repertoire features between control and disease cohorts. Overall, participants in the culture, infection, and cancer groups showed reduced IGH and TRB diversity relative to controls for the entire range of diversity measures (Fig. 3a; Supplementary Fig. 8a). Individuals with cancer generally had higher T cell:B cell (T:B) ratios, whereas individuals with infections displayed lower T:B ratios, consistent with the known roles of these cell types in these conditions (Fig. 3b; Supplementary Fig. 9). Specifically, the median *D*^1^ was 17,109 in non-control groups vs. 23,243 in controls, indicating fewer small clones in disease groups. Conversely, the median *D*^∞^ was 40 in non-control groups compared with 61 in controls, showing that the dominant clone is larger in disease groups. These observations confirm earlier reports and demonstrate a statistically significant trend toward clonal expansion in disease groups^14,31–33^.

**Figure 3.**
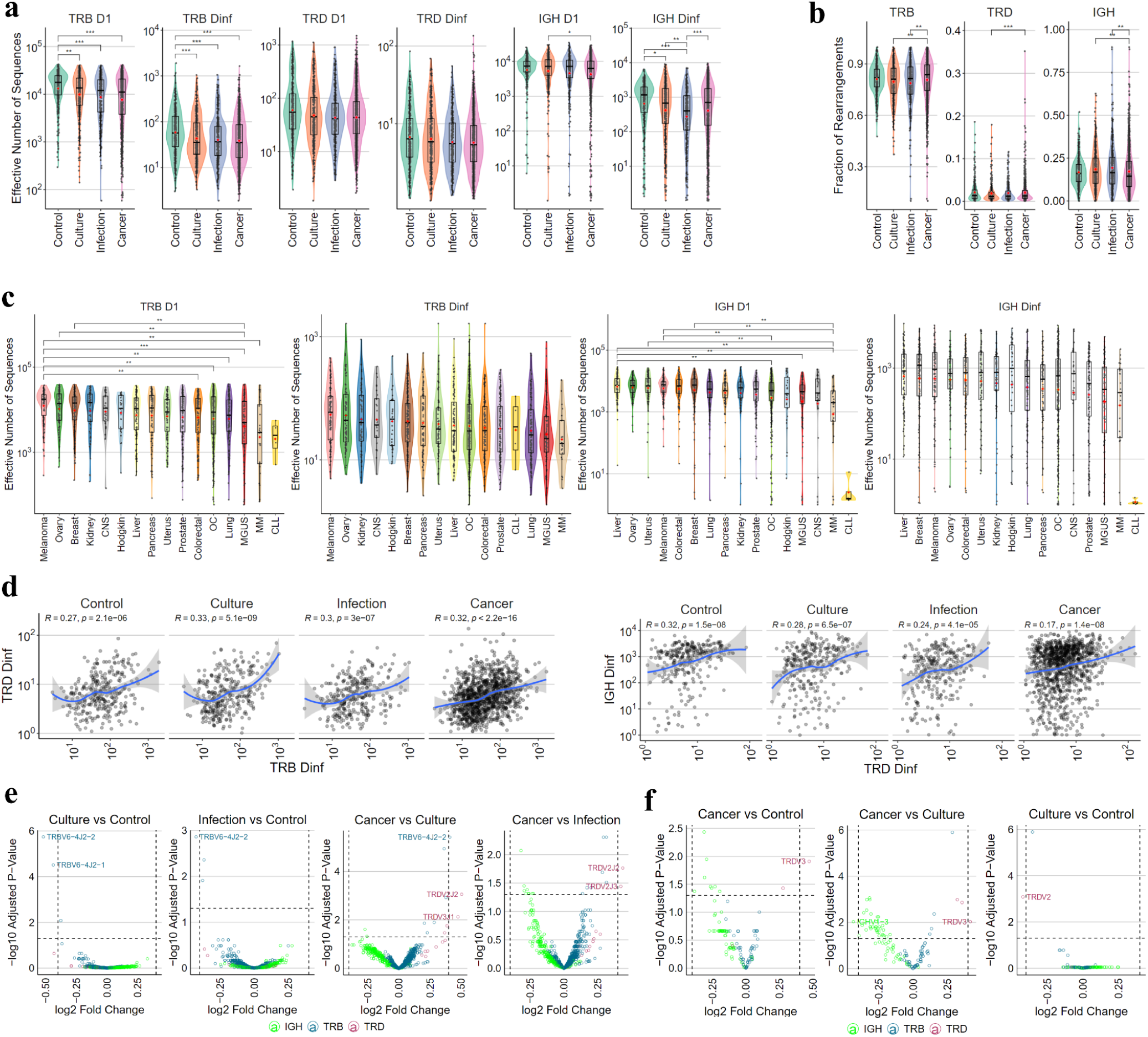
Immune repertoire analysis of control and disease groups. **a)** *D*^1^ and *D*^∞^ of different chain types in each group. **b)** Fraction of chain types in each group. **c)** *D*^1^ and *D*^∞^ for TRB and IGH chains in cancer subtypes. Only p values <0.01 are shown. **d)** Spearman correlation of Hill order infinity for TRD with TRB and IGH. **e-f)** Differential gene usage analysis between groups that demonstrated genes with differential usage for VJ pairs (e) or V gene (f). CLL, chronic lymphocytic leukemia; MM, multiple myeloma; CNS, central nervous system; MGUS, monoclonal gammopathy of undetermined significance; OC, other cancer. In box and violin plots statistical test was performed using a robust linear regression model (with covariate adjustment) and p values are FDR or Benjamini-Hochberg adjusted. * represents p value<0.05; ** p<0.01; *** p<0.001. Gene usage analysis was performed using limma-voom (with covariate adjustment) and p values are Benjamini-Hochberg adjusted.

The large size and diverse composition of the HuBIE cohort permitted intra–cohort comparisons, uncovering distinct immunological–diversity patterns among cancer subtypes and, for bloodstream infections, across different bacterial pathogens (Fig. 3c; Supplementary Fig. 8b–c). Both multiple myeloma (MM) and chronic lymphocytic leukemia (CLL), two hematologic malignancies, exhibited markedly reduced IGH diversity from *D*^1^ to *D*^∞^. A comparable reduction in IGH diversity was also seen in monoclonal gammopathy of undetermined significance (MGUS), the premalignant precursor to MM. Participants with CLL displayed the lowest IGH *D*^∞^ of all, consistent with dominance of a single large malignant clone. In contrast, Hodgkin lymphoma, also a hematologic malignancy, showed IGH diversity comparable to that of participants with solid tumors. The cohort also enabled simultaneous analysis of TCR subsets (Fig. 3c; Supplementary Fig. 8b-c). This analysis uncovered markedly low TRB *D*^1^ diversity in MGUS, MM, and CLL participants, indicating a reduced overall clone count. A very low TRB *D*^∞^, reflecting large clonal expansion of a single clone, was observed in MGUS and MM but not in CLL. These findings illustrate the hypothesis–generating power of large–scale immune–repertoire studies. Conversely, participants with bloodstream infections displayed a broadly uniform immunological–diversity profile, regardless of the bacterial pathogen isolated.

Immune–repertoire diversity showed correlated patterns across the various chain types. TRB, and to a lesser extent IGH, were positively correlated with TRD across the full range of diversity measures (*D*^0^–*D*^∞^; Fig. 3d; Supplementary Fig. 10). For *D*^0^, TRB and IGH displayed a negative correlation with TRD, likely reflecting a down–sampling artifact because the non–downsampled data did not show the negative correlation while retained a positive TRB/IGH–TRD correlation (Supplementary Fig. 11). CDR3 charge and length did not differ significantly among conditions, except for a few specific cancer subtypes (Supplementary Fig. 12).

The enormous combinatorial space of antibody and T–cell–receptor (TCR) sequences leads to low pairwise sequence overlap between individuals^22,34^. HuBIE’s size and breadth offered a unique opportunity to examine this phenomenon at scale. We calculated immune–repertoire overlap for every participant pair and stratified the values by group. Participants in the cancer cohort showed significantly lower IGH, TRB, and TRD overlap with one another than control participants (Supplementary Figs. 3c & 4b), a pattern consistent with cancer imposing divergent selection pressures on the adaptive immune system. We then performed a differential gene–usage analysis to identify differences in V, J, and VJ–combination usage between groups (Fig. 3e–f; Supplementary Figs. 13 & 14). Overall, gene–usage patterns were similar across the main groups; only a handful of TRB VJ pairs were under–used in infection and culture cases, whereas a few TRD and TRB VJ pairs were over–used in cancer relative to culture and infection. Specifically, TRBV6–4J2–2 was less frequent in culture and infection samples but more frequent in cancer. Likewise, a small set of IGH V genes showed reduced usage, while several TRD V genes were more frequently used in cancer. Nevertheless, specific cancer subtypes—especially hematologic malignancies such as CLL, MM, MGUS, and Hodgkin lymphoma—exhibited many significant gene–usage differences versus controls, with numerous IGH V, J, and VJ pairs being used less frequently (Supplementary Fig. 13).

### Temporal Trends and Survival Status

We analyzed longitudinal samples from 419 HuBIE participants to characterize immune–repertoire dynamics during acute illness. First, we confirmed that high–frequency clonotypes were reproducibly quantified (see Methods and Supplementary Fig. 16). As a case study, we focused on a single participant with a confirmed infection. This individual contributed 12 longitudinal samples spanning their entire illness (Fig. 4a). The participant was first diagnosed with a coagulase–negative Staphylococcus (CoNS) infection, which was superseded by a *Staphylococcus aureus* (SAU) infection beginning at time–point 2 (39 days after the initial sample) and lasting until time–point 8 (81 days after the initial sample). The combined frequency of the top 5 TRB and IGH clones initially declined, then rose to a peak around time–point 7–8. The subsequent decline in clonotype frequency coincided with infection resolution. Thus, the expansion of these dominant clones reflects a targeted immune response, and their contraction aligns temporally with pathogen clearance.

**Figure 4.**
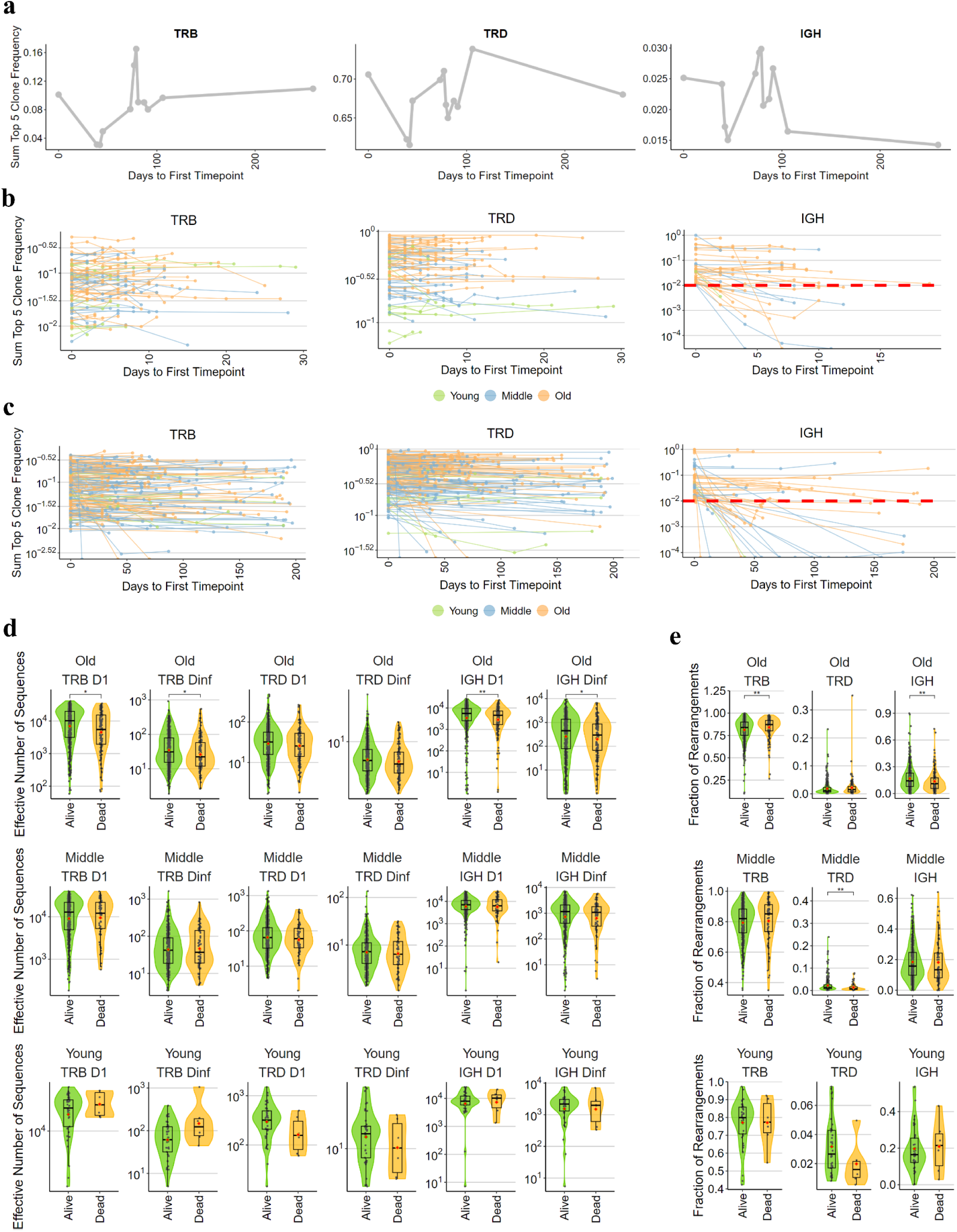
Immune repertoire analysis in longitudinal participants and with regards to survival status. **a)** Tracking the sum frequency of top 5 clonotypes in a participant with 12 timepoints. **b)** Tracking the frequency of top 5 clonotypes in longitudinal infection participants. **c)** Tracking the frequency of top 5 clonotypes in longitudinal cancer participants. **d)** Chain-level *D*^1^ and *D*^∞^ for cancer participants who were still alive and those who died in three age groups. **e)** TRB, TRD, and IGH chain fractions for cancer participants who were still alive and those who died in three age groups including young (<40 years old), middle age (40-65) and old (>65 years old). For comparison of dead and alive cases, statistical test was performed using a robust linear regression model (with covariate adjustment) and p values are FDR or Benjamini-Hochberg adjusted. * represents p value<0.05; ** p<0.01; *** p<0.001. Young (<40 years old). Middle-age (40-65 years old). Old (>65 years old). For clone tracking in infection and cancer participants, only samples with >40,000 rearrangements were considered, and for IGH only top five clones with sum frequency of >0.01 were included.

We tracked the longitudinal frequencies of the top TRB, TRD, and IGH clones in infection (≤30 days) and cancer (≤200 days) cohorts (Fig. 4b–c). Across most participants, the summed frequency of the five most abundant TRB and TRD clones remained essentially unchanged in both conditions. This stability likely reflects that most participants’ disease status/group membership remained stable across most time points. In contrast, top IGH clone frequencies were stable in most infection cases but tended to decline over time in many cancer participants. This opposite behavior likely stems from the distinct roles of B cells: during infection the leading IGH clones are actively engaged against the pathogen, maintaining their frequencies, whereas in cancer the dominant IGH clonotypes are possibly unrelated to the anti–tumor response, enabling their abundances to be more dynamic. Nevertheless, we identified a few infection cases in which initially high–frequency IGH clones fell sharply later on. For instance, in one infection case (MDS_11991) the combined frequency of the top 5 IGH clones dropped from 0.39 at the first visit and 0.29 at the second visit to undetectable by the third. This sample had 4 time points and the forth specimen was subsequently re–classified as a culture(-negative) sample collected 43 days after the initial draw, indicating that the infection was no longer detectable. This abrupt loss of dominant IGH clones aligns with the participant’s clinical outcome.

Immune diversity declines with age, implying that reduced diversity might increase mortality risk. Consistent with this, prior studies have shown that lower immune–repertoire diversity—especially in T–cell receptors—is associated with poorer overall survival in cancer participants^35,36^. Consequently, we examined whether immune diversity varies across age groups and could serve as a prognostic indicator of death. Among cancer participants at baseline, about 19 % died during follow–up. In the young (< 40 yr) and middle–aged (40–65 yr) cohorts, we observed no significant difference in repertoire diversity between survivors and non–survivors. By contrast, in the elderly (> 65 yr) cohort, deceased participants exhibited significantly lower diversity in both IGH and TRB repertoires (Fig. 4d; Supplementary Fig. 15a). Furthermore, the elderly cancer participants who died showed reduced IGH fractions together with elevated TRB fractions, which may reflect a late, ineffective surge in anti–cancer T–cell activity (Fig. 4e). Conversely, among middle–aged cancer participants who died, we detected lower TRD chain fractions and alterations in CDR3 characteristics such as length and charge (Supplementary Fig. 15b–c).

## Discussion

Immune–repertoire sequencing is attracting growing interest because of its potential for disease diagnosis and therapy. Many large–scale cohort projects infer immune–receptor repertoires from whole–transcriptome datasets (e.g., the Cancer Genome Atlas, TCGA), a strategy that can compromise resolution and specificity^37,38^. In contrast, dedicated immune–repertoire sequencing efforts usually profile only TCRs or only BCRs and are typically limited by modest cohort sizes or a lack of demographic diversity. To address these gaps, we created the Human Blood Immunome Encyclopedia (HuBIE). To our knowledge, HuBIE is the largest multi–disease cohort that captures simultaneous TCR and BCR repertoire data together with extensive clinical metadata. The cohort constitutes a landmark resource for probing how immune–repertoire dynamics interact with demographic variables and disease phenotypes. Integrating large–scale TRB, TRD, and IGH repertoire data from diverse populations allowed us to uncover new insights into the ways age, sex, race, and underlying pathology sculpt adaptive immunity.

Our assessment of immune–repertoire diversity conforms with earlier work that reported reduced repertoire richness and overall diversity in diseased specimens, a pattern interpreted as clonal expansion of particular cell populations^14,31–33^. In our dataset, the effect was especially pronounced: TRB clonotypes showed marked clonal expansion in cancer cases, whereas IGH clonotypes expanded in infection–related disorders compared with controls. These results reinforce the prevailing view of the adaptive immune system mounting distinct responses to pathogenic versus malignant challenges.

Importantly, different cancer and infection subtypes displayed separate repertoire signatures. Cancer participants, especially individuals with hematological malignancies such as chronic lymphocytic leukemia (CLL) and monoclonal gammopathy of undetermined significance (MGUS), exhibited marked hypodiversity in both IGH and TRB repertoires, indicative of clonal expansion of malignant cells.

Hodgkin lymphoma stood apart from other hematological cancers (possibly because Reed–Sternberg cells do not circulate, which may produce a muted systemic immune response to the tumor). Conversely, bloodstream infections maintained overall repertoire diversity even though particular IGH clones expanded, suggesting pathogen–driven selection pressures. The observed negative correlation between TRB and IGH diversity, together with the positive coupling of TRB and TRD diversities, underscores the intricate regulatory networks that operate within the adaptive immune system.

Patient age proved to be the primary factor driving repertoire decline. IGH diversity decreased sharply after age 60, a pattern that reflects increased clonality. In contrast, TRB diversity declined more gradually with age, suggesting intrinsic differences in T–cell homeostatic mechanisms. TRD diversity showed a steady age–related decline, consistent with the known reduction in γδ T–cell production^27,29^. Collectively, these trends indicate that aging alters selection pressures, with IGH diversity appearing especially vulnerable to stochastic or environmental factors. Racial differences in repertoire architecture were pronounced: Asian participants showed higher IGH diversity and lower TRB diversity than white participants, while Black and Hispanic individuals displayed intermediate values. These disparities also encompassed gene–segment usage and chain–type frequencies, highlighting the combined influence of genetic background and environmental exposures on shaping adaptive immunity. Such observations have potential implications for personalized medicine, because repertoire variation may affect vaccine responsiveness and autoimmune susceptibility. The wide demographic coverage of the cohort also permitted novel analyses of inter–individual repertoire overlap. Cancer participants exhibited reduced immune–repertoire sequence similarity relative to other groups, possibly reflecting more divergent antigenic landscapes. Conversely, younger individuals displayed greater sequence similarity across participants, particularly for the TRD chain. The age–associated changes in TRD appear to stem from altered gene–segment usage rather than a uniform loss of sequences, indicating a biased expansion of particular clones that could influence immune memory.

Prior work has shown that greater immune–repertoire diversity—especially within T–cell compartments—is linked to improved survival among cancer participants^35,36^. Our results are consistent with those reports: cancer participants who survived exhibited higher overall immune–repertoire diversity than participants who died, a difference that was most pronounced in the TRB chain.

Importantly, the diversity advantage reached statistical significance only in the elderly subgroup. Conversely, among middle–aged cancer participants we detected differences in CDR3 characteristics—specifically length and net charge—between survivors and non–survivors. Taken together, these findings imply that overall repertoire diversity serves as a more informative prognostic factor in older participants, whereas finer–grained attributes such as CDR3 physicochemical properties may provide superior prognostic insight in younger individuals.

Although the size and demographic breadth of HuBIE afford novel insights, the investigation is subject to several limitations. First, it is limited by its observational nature and by reliance on self–reported demographic information. Future work should seek to replicate these observations in prospective longitudinal cohorts of real-world participants presenting for care and to elucidate the functional consequences of the identified repertoire variations. Second, full-length and paired-chain sequence was not performed as it is not yet economically practical at this scale. Third, because samples were collected only relatively recently, we do not yet have long-term followup data on most participants; however, our data provides a baseline for followup as time goes on. Overall, HuBIE provides a valuable framework for dissecting how biological and environmental factors shape immune diversity, with direct relevance to precision immunology, as well as a large and richly characterized, freely available dataset for a range of future analyses, including disease classification, regression studies, and discovery.

## Materials and Methods

Work was approved by Beth Israel Deaconess Medical Center (BIDMC)’s institutional review board (BIDMC IRB 2021P000899).

### Sample collection and DNA extraction

Diagnostic blood samples discarded in BD Vacutainer® EDTA tubes (Becton, Dickinson and Company, Franklin Lakes, USA) were obtained from participants at Beth Israel Deaconess Medical Center (BIDMC) between 2022 and 2024. Longitudinal collections from multiple participants yielded a total of 2,614 blood samples representing 1,941 individuals targeted for the study. We focused on participants with common cancers, rare cancers, and those with confirmed or suspected bacterial bloodstream infections. Additional discarded blood from participants without these conditions and without known immune–system–affecting disorders served as controls. Details of the sample–selection criteria are provided in Supplementary Tables 2 and 3.

We collected 2 mL of blood discards within 10 hours of drawing and isolated peripheral blood mononuclear cells (PBMCs) using SepMate™-15 (IVD) tubes (STEMCELL Technologies, Vancouver, Canada) with Ficoll-Paque PLUS density gradient medium (Cytiva, Little Chalfont, United Kingdom), following the SepMate protocol. PBMCs were cryopreserved in CELLBANKER 2 (AMSBIO, Oxfordshire, UK) at -80°C. Frozen PBMCs were thawed in a 37°C water bath, washed with MACS® Separation Buffer (Miltenyi Biotec, Bergisch Gladbach, Germany), and a small portion of the cell suspension was used for differential cell counting on an Automated Hematology Analyzer XN-10™ (Sysmex Corporation, Kobe, Japan) to determine lymphocyte ratios in each PBMC sample. The remaining PBMCs were lysed with Proteinase K and RNase A (both from MACHEREY-NAGEL, Düren, Germany), followed by the addition of Lysis Buffer BQ1 from the Blood QuickPure kit (MACHEREY-NAGEL). The lysates were then stored at -20°C until DNA extraction. DNA was extracted from PBMCs using the NucleoSpin 96 Blood QuickPure, 96-well kit (MACHEREY-NAGEL) on a liquid handler Biomek i5 (Beckman Coulter Life Sciences, Indianapolis, USA). The final DNA elution was performed using 10mM Tris-HCl buffer (pH 8.0).

To prepare DNA templates for the immunoPETE method, DNA sample concentrations were adjusted with 10 mM Tris-HCl buffer (pH 8.0) to achieve the recommended 500 ng of target DNA (lymphocyte DNA) per reaction, based on PBMC differential cell count results. Longitudinal samples from the same participant were placed on the same immunoPETE 96-well plate. Duplicates or triplicates of each sample were included on the plate along with 3 negative and 3 positive controls. The positive control was prepared by pooling DNA extracted from PBMCs of a healthy individual (Human Peripheral Blood Mononuclear Cells, Frozen, #70025, STEMCELL Technologies, Vancouver, Canada) at 495 ng of lymphocyte DNA per immunoPETE run with 5 ng of DNA from the B-cell line ARH-77 (American Type Culture Collection). This DNA mixture was prepared in a single batch, and aliquots of the batch were used as the positive control in each immunoPETE run to ensure consistency across plates. Under these conditions, samples from different health conditions, sampling dates, and other categories were intentionally distributed across various immunoPETE runs to reduce potential batch effects. Totally 7,427 immunoPETE products underwent Next-generation sequencing with unique molecular identifiers.

### immunoPETE sequencing

Immunoglobulin heavy chains for BCR (IGH) and both beta and delta chains for TCR (TRB and TRD) were captured and sequenced using immunoPETE technology from Roche. The immunoPETE workflow was automated on a liquid handler Sciclone G3 NGSx iQ Workstation (CLS152236, Revvity Germany GmbH). Briefly, from genomic DNA, an initial primer extension was performed using KAPA LongRange HotStart ReadyMix with dye (KK3602, Sigma Aldrich) and a pool of 173 V-gene oligonucleotides (Integrated DNA Technologies Germany, GmbH) containing a unique molecular identifier (UMI) sequence and a universal amplification sequence. An exonuclease I (M0568L, New England Biolabs) treatment and bead-based purification using KAPA HyperPure Beads were then performed to remove remaining oligonucleotides and unextended genomic DNA. Next, a master-mix of KAPA LongRange HotStart ReadyMix with dye (KK3602, Sigma Aldrich) and a pool of 25 J-gene oligonucleotides (Integrated DNA Technologies Germany, GmbH), an i7-bridging primer (Integrated DNA Technologies Germany, GmbH) and i7/i5-sequencing primers with dual unique indexes (KAPA UDI Primer Mixes) were added to the purified V-gene-primed templates for J-gene primer extension.

Resulting libraries were purified using KAPA HyperPure Beads before quantification and fragment analysis. 144 libraries (equivalent to 1.5 96-well microtiter plates and containing at least 3 positive and 3 negative controls) were pooled in equal volume to create a library pool before another round of bead-based purification using KAPA HyperPure Beads, quantification and fragment analysis. Four library pools were sequenced in each of the 4 lanes of an S4 flow cell on an Illumina NovaSeq 6000 platform using 2×150-base paired-end reads. All library preparation and sequencing were performed at Signature Diagnostics GmbH / Roche Potsdam, Germany.

### immunoPETE data processing

V/J-gene annotations and CDR3 sequences were extracted from the raw sequencing data using a bioinformatics pipeline designed by Roche to process immunoPETE data. This involved validating UMI patterns, down-sampling raw FASTQ sequences to a maximum of 50 million valid reads per library, trimming primer templates from read mate pairs, and aligning them to IG and TR references from the IMGT database. Subsequently, gene-aligned reads with a quality score greater than Q30 were filtered, and 5 million on-target reads were subsampled and clustered into UMI families based on their CDR3 content and UMI sequence.

### Data downsampling and calculation of Hill numbers

For comparisons of metrics affected by cell count such as Hill numbers we randomly downsampled results of each sample to 50k functional or productive rearrangement/cell count with 25 bootstraps (Supplementary Fig. 1). This resulted in downsampling of 93.3% of the samples, we also removed samples with low cell count (≤40K). For each downsampled boot strap we calculated each metric and used the average of measured metrics from all the bootstraps for each sample as the final value. For the pairwise immune repertoire overlap, due to computation time, we only used the first downsampled bootstrap. D numbers can be calculated using the Greylock Python package^39^. To explore immune repertoire richness and diversity we calculated D numbers at orders *D*^0^, *D*^1^, *D*^2^, and *D*^∞^ as follows.

*D*^0^ represents the richness or number of unique clones. *D*^1^ represents the exponential of the Shannon entropy and highlights the effective number of common species. *D*^1^ is calculated using the *e*(H) formula where H is Shannon entropy and calculated as –𝚺(*p_i_*ln(*p_i_*) where 𝑝ᵢ represents the proportion of the i^th^ species/clone. *D*^2^ represents the inverse of the Simpson index and highlights the dominant species. We calculated *D*^2^ using the 1/Σ𝑝ᵢ² equation where 𝑝ᵢ represents the proportion of the *i*^th^ species/clone. *D*^∞^ represents Berger-Parker index for clonality which is the inverse proportion of the most frequent species/clone 1/max(*p*_i_).

### Calculation of the CDR3 net charge

We calculate the net charge of CDR3 based on Henderson-Hasselbalch equation at pH 7.4. We used the following equation:

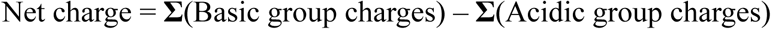

For basic groups charge including N-terminus, Lys, Arg, His the following equation was used:

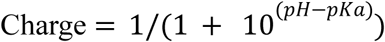

For acidic groups charge including C-terminus, Asp, Glu, Cys, Tyr the following equation was used:

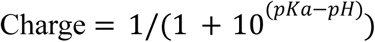

The pKa values for the ionizable groups captured from Lehninger’s Principles of Biochemistry 8th edition.

### Immune repertoire pairwise overlap analysis

We calculated pairwise overlap between two samples using a Sorensen-Dice coefficient like equation:

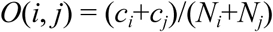

Where *O*(*i*, *j*) is the overlap score between two samples. *c_i_* is the number of functional CDR3 rearrangements in the first sample with the same V and J genes and the maximum hamming distance of 1 with the second sample. *c_j_*is the number of functional CDR3 rearrangements in the second sample with the same V and J genes and the maximum hamming distance of one with the first sample. *N_i_* is the number of all functional CDR3 rearrangements in the first sample and *N_j_* is the number of all functional CDR3 rearrangements in the second sample.

### Clone-tracking reproducibility

To examine the possibility of clone tracking we examined the overlap through exact match (number of common rearrangements / all rearrangements, in a pairwise comparison) between replicate samples from the same participant in 10 randomly selected participants with cancer or infection with multiple timepoints. As anticipated, hierarchical clustering of immune repertoires overlap showed that different time points from the same participant and replicates from the same sample were consistently clustered together (Supplementary Fig. 16a). This was especially evident for the TRB, which constitutes a significant portion of the immune repertoire. We then evaluated the consistency of the frequency of the most abundant clonotypes across 100 randomly selected samples with three replicates. We found that the frequencies of the top clonotypes were highly consistent for TRB and, to a lesser extent, for the most frequent TRD clonotype (Supplementary Fig. 16b-d). When we aggregated the frequencies of the top five clonotypes, their summed frequency was almost perfectly consistent between replicates for both TRB and TRD, but not for the IGH (Supplementary Fig. 16e). However, for IGH, when the frequency was greater than 0.01 in the first replicate, then they mostly showed consistent frequency over other replicates (Supplementary Fig. 16f). This finding suggests that the aggregated frequency of the most abundant clones is a reliable metric for tracking changes over time for TRB and TRD, and for IGH only if the frequency is greater than 0.01 and the higher the more reliable.

## Author contributions

Conceptualization: RA, HM, FR, HA, FR. Methodology: RA, HM, HA, FR. Software: AM, HA, HZ, VA, CT. Validation: AM, MY, EC, JAB, DB, DS, TS, SD, SR, SM, SMN, SS, US. Formal Analysis: VA, AM, RA, CT, SMN, ZGD, HZ. Investigation: AM, MY, EC, JAB, DB, DS, TS, SS, US. Resources: SD, SR, SM, DB, US. Data Curation: AM, RA, CT, SMN, ZGD. Writing – Original Draft: VA, RA. Writing – Review & Editing: RA, VA, HM, AM. Visualization: VA, RA. Supervision: RA, HM, MY, US, DK. Project Administration: RA, HM, DK. Funding Acquisition: RA, SU.

