## Supplementary Table 2 for "HuBIE: The Human Blood Immunome Encyclopedia Of TCRs and BCRs in Bloodstream Infections and Cancer"

**Supplementary Table 2: Sample categories by participants' conditions**

| Group | Description | |
|---|---|---|
| **Cancers** | The participant's chart contained an ICD-10 code within the prior year indicating any of cancer listed below or participants with a protein electrophoresis result <10 days indicating abnormal gamma bands, suggesting a possible monoclonal gammopathy. | |
| | breast | lung |
| | central nervous system | melanoma |
| | colorectal | ovary |
| | Hodgkin lymphoma | pancreas |
| | kidney | prostate |
| | liver | uterus |
| | leukemia | monoclonal gammopathy |
| **Bacterial blood stream infection** | Participants with a recent (within the prior 3 days) blood sample was positive for an infection when cultured in the clinical microbiology lab. | |
| | Coagulase-negative Staphylococci (CoNS) | *Pseudomonas aeruginosa* |
| | Enterobacterales | *Staphylococcus aureus* |
| | Enterococci | Viridans streptococci |
| **Blood culture controls** | A recent (within the prior 3 days) blood sample was negative for any infection when cultured in the clinical microbiology lab | |
| **Controls** | Blood is discarded from participants who were not in categories above and who did not have conditions below. They were collected matching the target population in gender, race, and age distribution. | |
| | HIV | Crohn's disease |
| | lymphoma | Celiac disease |
| | History of transplant | Autoimmune hepatitis |
| | any type of cancer | Vitiligo |
| | obesity | Lupus |
| | autoimmune hemolytic anemia | Reactive arthropathy |

| | |
|---|---|
| Sarcoidosis | Juvenile arthritis |
| Type I diabetes | Sjögren syndrome |
| Autoimmune polyglandular failure | Ankylosing spondylitis |
| Autoimmune thyroiditis | Myasthenia gravis |
| Addison’s disease | Rheumatism |
| Amyloidosis | Fibromyalgia |
| Multiple sclerosis | other autoimmune disorders |
| moderate or severe asthma | liver or kidney failure |
