## Supplementary Table 3 for "HuBIE: The Human Blood Immunome Encyclopedia Of TCRs and BCRs in Bloodstream Infections and Cancer"

**Supplementary Table 3:** Additional participants metadata.

| Information | Description |
|---|---|
| **Race** | Self-reported race and/or ethnicity of the participant: White, Hispanic, Black/African-American, Asian or Pacific Islander, Native American, or unknown/not reported/not classified |
| **Sex** | Gender of the participant according to the medical record as of the blood sample date. |
| **Age** | Age of participant at time of blood sample; or "90+" whenever the participant was at least 90 years old. |
| **Antibiotics** | List of medications started before the blood sample date, and either current at the time of the blood sample or discontinued less than 30 days before the blood sample date; number of days given is the number of days before the blood sample date that the medication was started. |
| **Cancer medication** | List of medications started before the blood sample date, and either current at the time of the blood sample or discontinued less than 30 days before the blood sample date; number of days given is the number of days before the blood sample date that the medication was started. |
| **Sample dates** | All sample dates were encoded to protect participants' health information. |
| **Deceased date** | Whether there is a record in the database that the participant has died. |
