## Supplementary Figures for "HuBIE: The Human Blood Immunome Encyclopedia Of TCRs and BCRs in Bloodstream Infections and Cancer"

* Equal Contribution

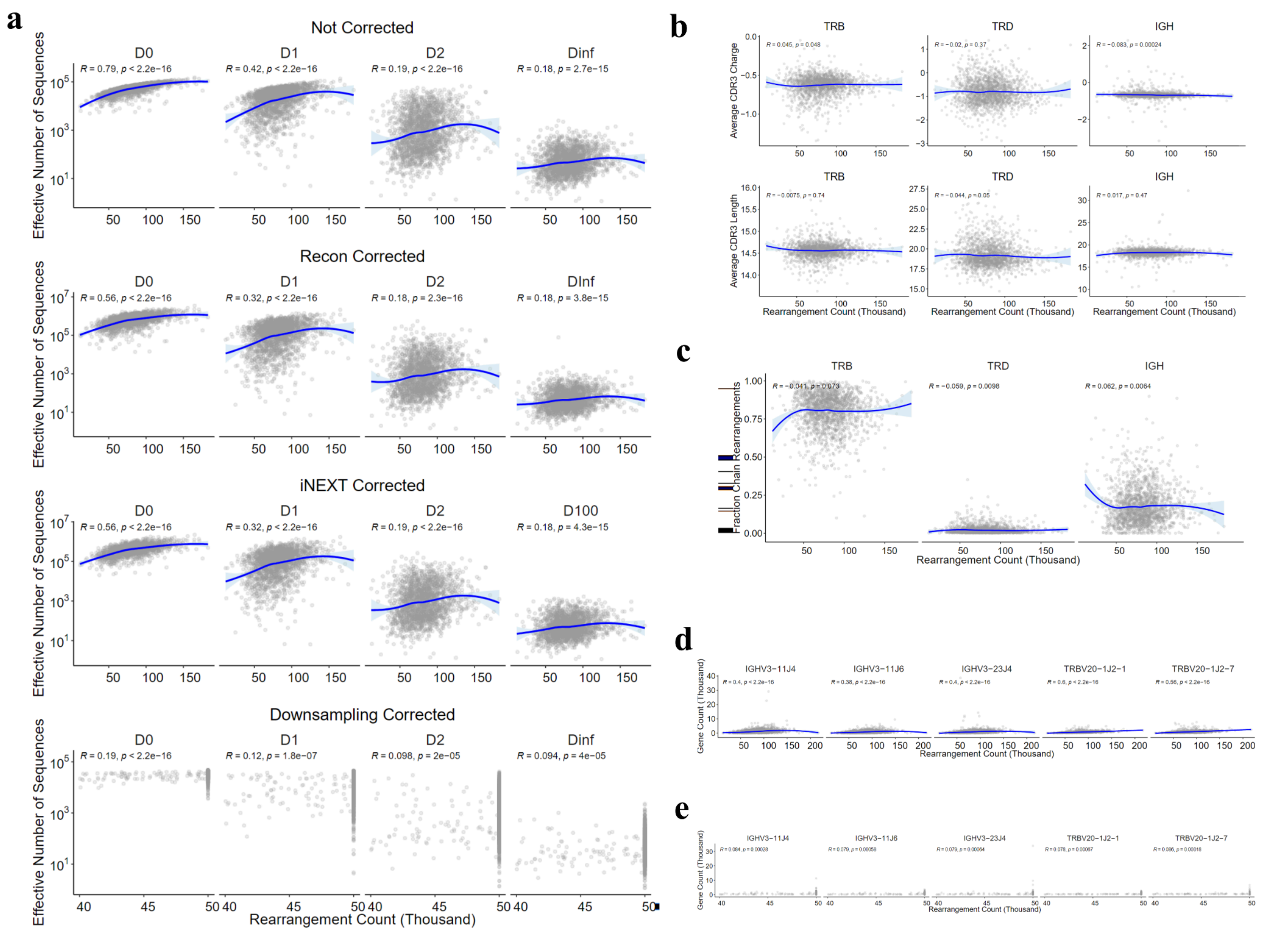


**Supplementary Figure 1.** a) Spearman correlation of $D^q$ numbers vs. cell count from un-corrected data (i.e. no downsampling) and data with sampling bias addressed using the iNEXT R package, Recon, and downsampling. Samples with less than 40,000 cells are also excluded after downsampling. **b**) Spearman correlation of cell count and average CDR3 length and charge from non-downsampled samples. **c**) Spearman correlation of cell count and TRB/TRD/IGH chain fractions calculated from non-downsampled samples. d) Spearman correlation of cell count and top gene pairs gene count from non-downsampled samples. e) Spearman correlation of cell count and the top gene pairs gene count calculated from downsampled samples.

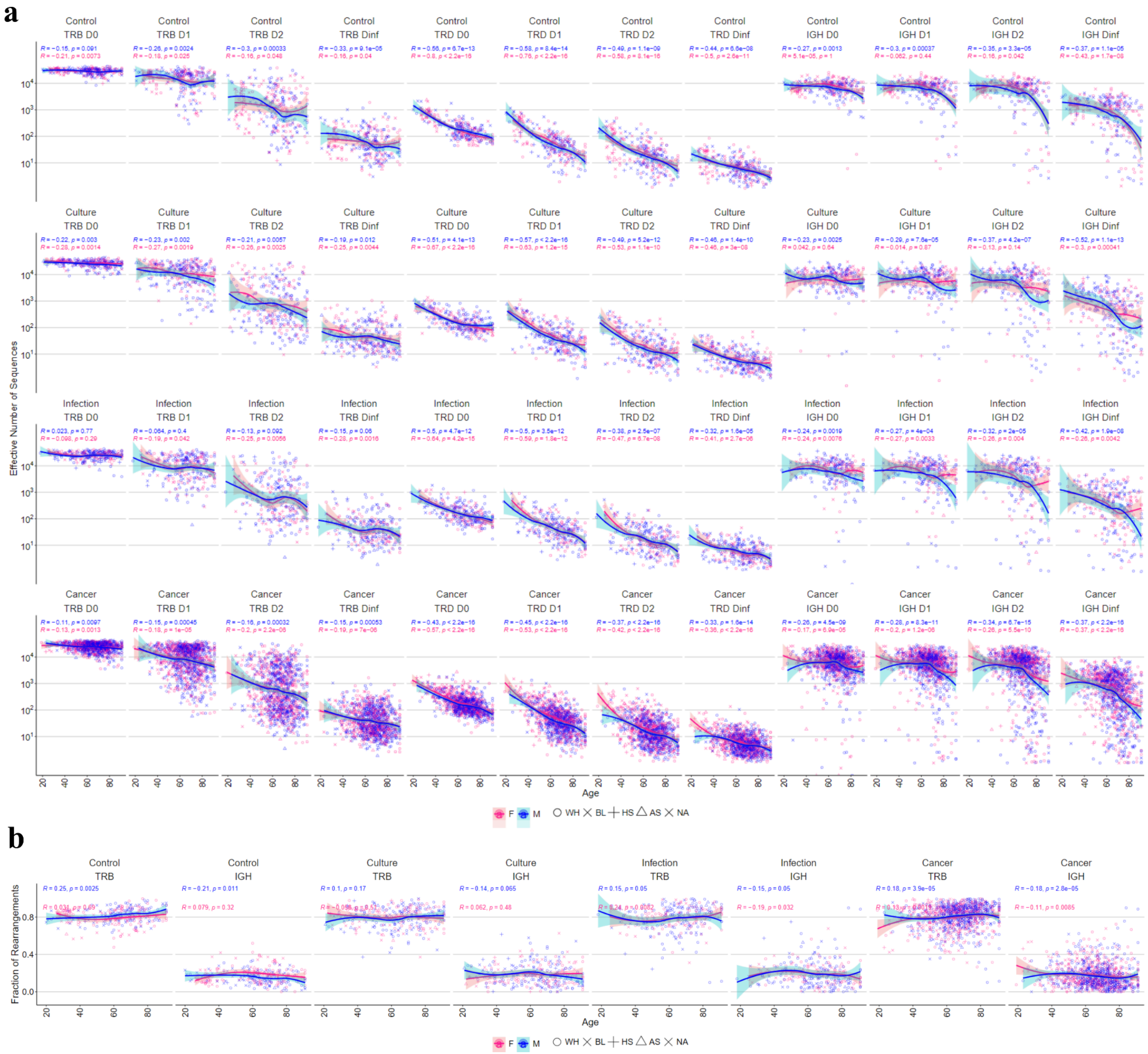


**Supplementary Figure 2**. a) Chain-level Spearman correlation of $D^q$ numbers with age in each group. b) Correlation of chain fraction for IGH and TRB with age in each group.

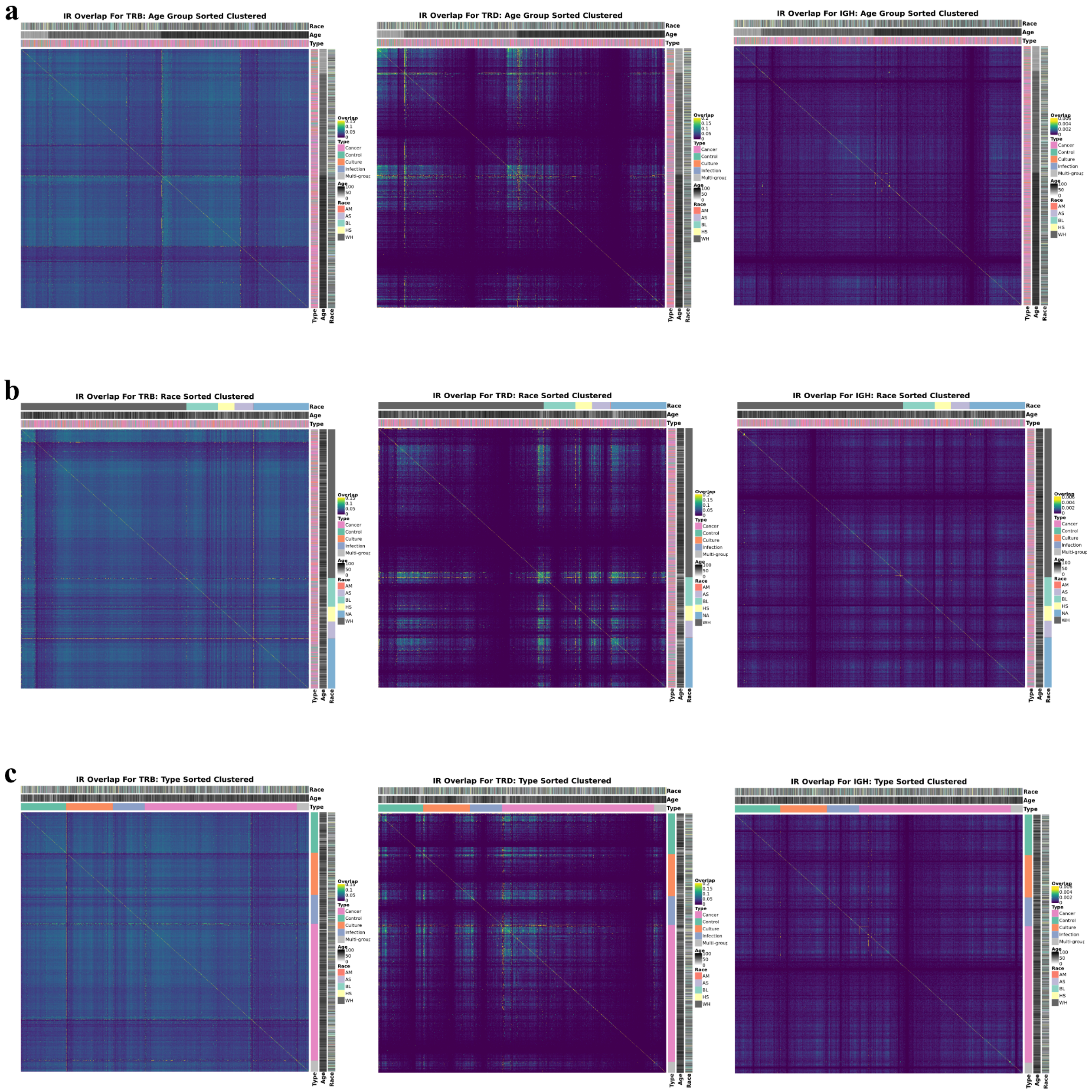


**Supplementary Figure 3.** Pairwise immune repertoire overlap. Heatmaps are sorted in different ways, age group sorted (a), race sorted (b), disease status or type sorted (c), and then clustering performed inside each category that sorting is performed based on that.

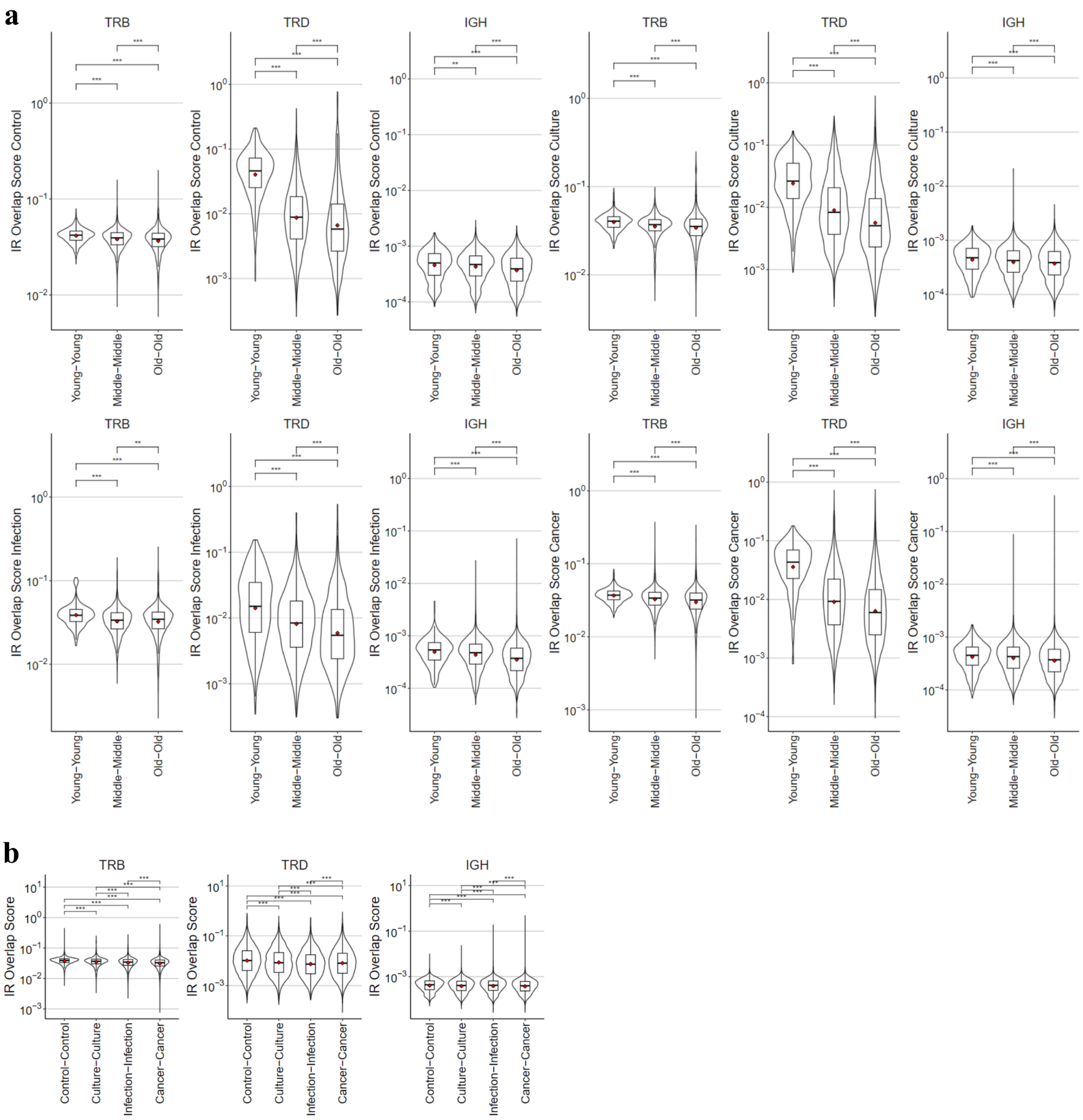


**Supplementary Figure 4.** Comparison of pairwise immune repertoire overlaps based on age groups (a) and disease groups (b). Wilcoxon statistical test was performed and p values are FDR or Benjamini-Hochberg adjusted. * represents p value<0.05; ** p value <0.01; *** p value <0.001. For gene usage plots p value is Benjamini-Hochberg adjusted.

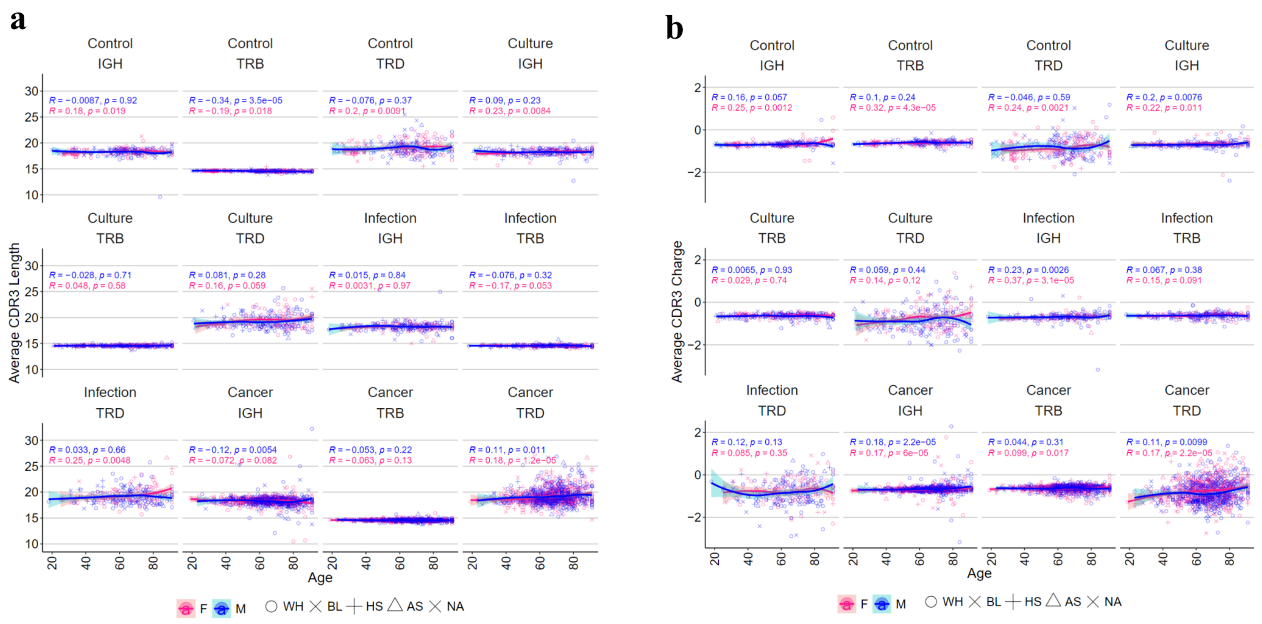


**Supplementary Figure 5.** Analysis of age and CDR3 charge and length with age. a) Spearman correlation of average CDR3 length with age in each condition. b) Spearman correlation of average CDR3 charge with age in each condition.

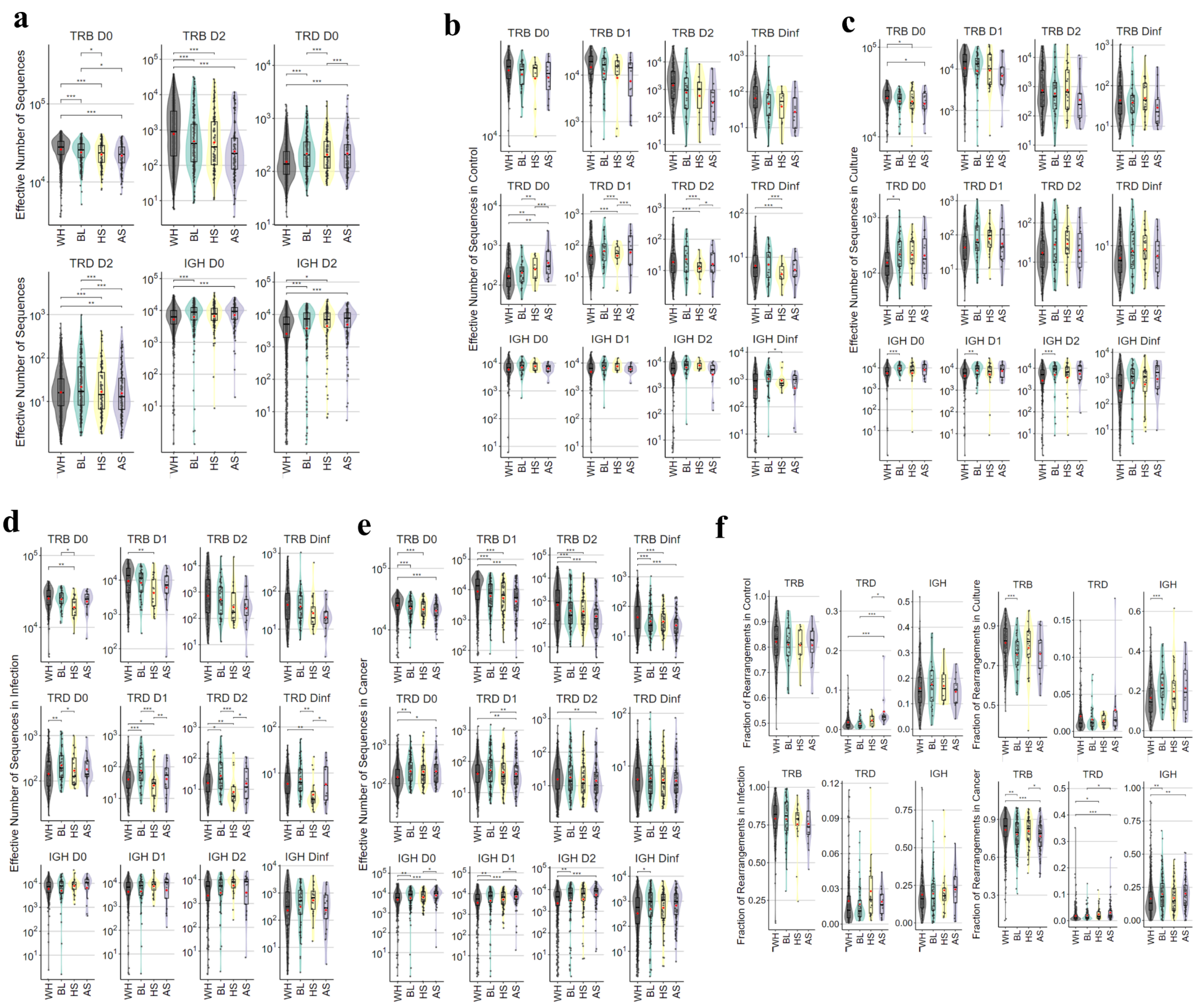


**Supplementary Figure 6** . Analysis of race and immune repertoire diversity. a) Immune repertoire $D^q$ numbers at orders 0 and 1 for different race groups across the cohort. b-e) $D^q$ numbers in different races in each control (b), culture (c), infection (d), and cancer (e) groups. f) Fraction of TRB, TRD, and IGH chain in different races in each condition. WH, white; BL, black; HS, hispanic; AS, asian. Statistical test was performed using a robust linear regression model (with covariate adjustment) and p values are FDR or Benjamini-Hochberg adjusted. * represents p value<0.05; ** p<0.01; *** p<0.001. native Americans (AM) are excluded as there were very few AM samples.

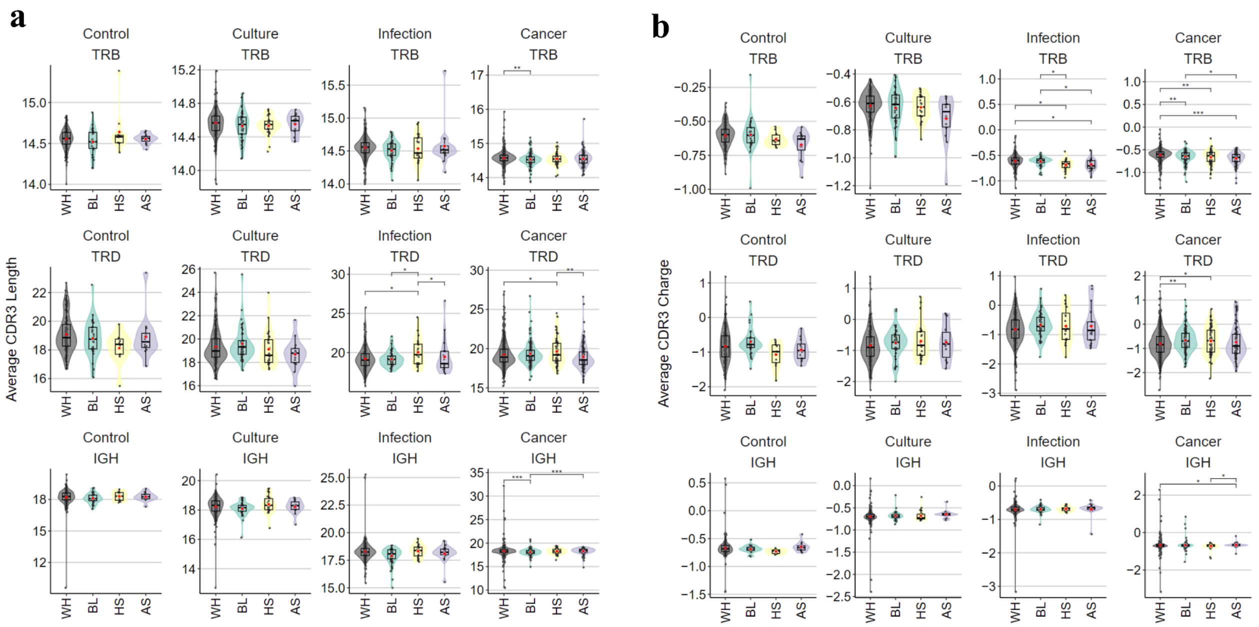


**Supplementary Figure 7**. Analysis of race and CDR3 length and charge. a) Average CDR3 length in each race in each condition. b) Average CDR3 charge in each race in each condition. WH, white; BL, black; HS, hispanic; AS, asian. Statistical test was performed using a robust linear regression model (with covariate adjustment) and p values are FDR or Benjamini-Hochberg adjusted. * represents p value<0.05; ** p<0.01; *** p<0.001. native Americans (AM) are excluded as there were very few AM samples.

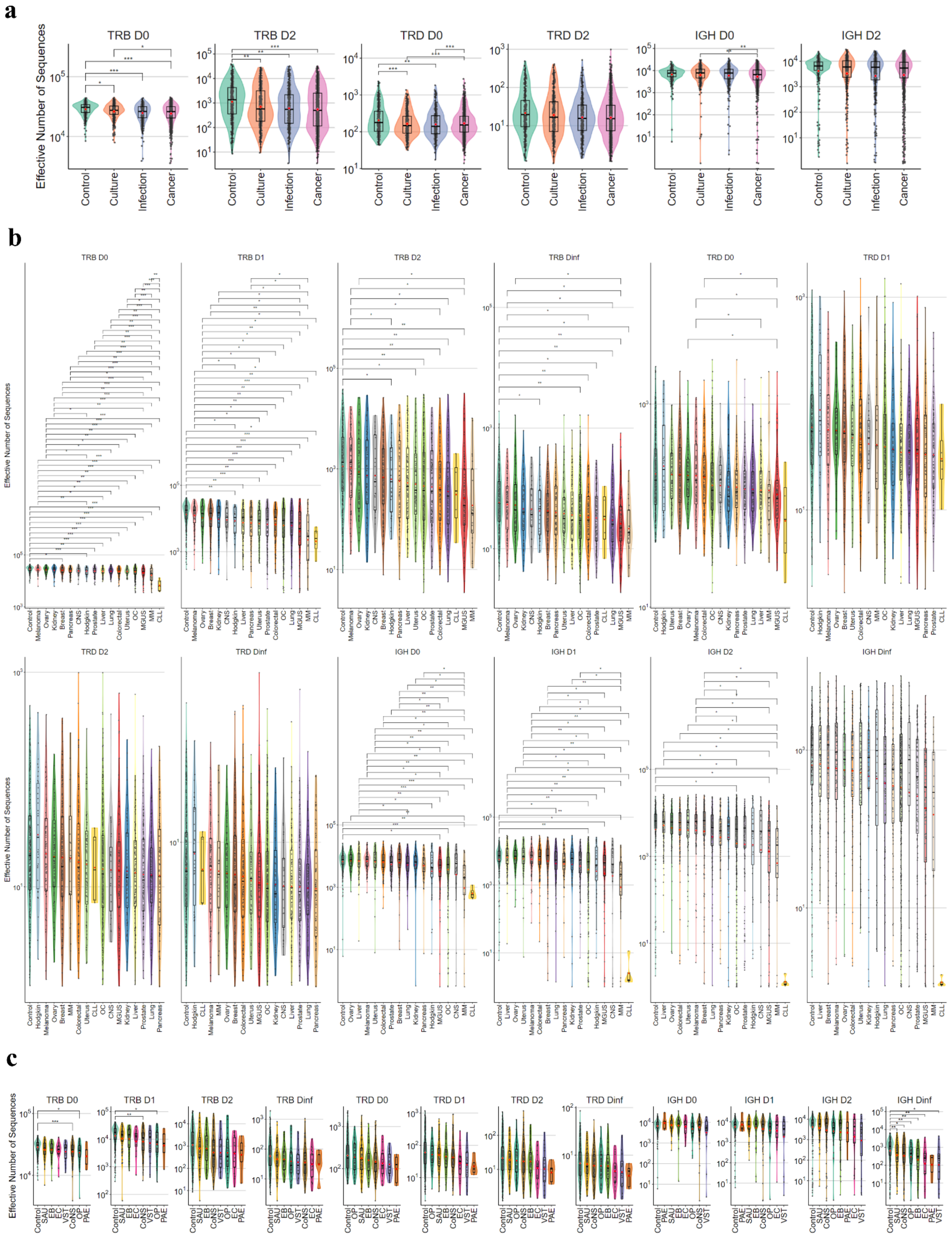


**Supplementary Figure 8**. Analysis of immune repertoire diversity based on disease status. a) $D^q$

numbers at orders 0 and 1 in each condition. b) $D^q$ numbers for cancer subtypes and their comparison to control and each other. c) $D^q$ numbers for infection subtypes and their comparison to control and each other. Statistical test was performed using a robust linear regression model (with covariate adjustment) and p values are FDR or Benjamini-Hochberg adjusted. * represents p value<0.05; ** p<0.01; *** p<0.001.

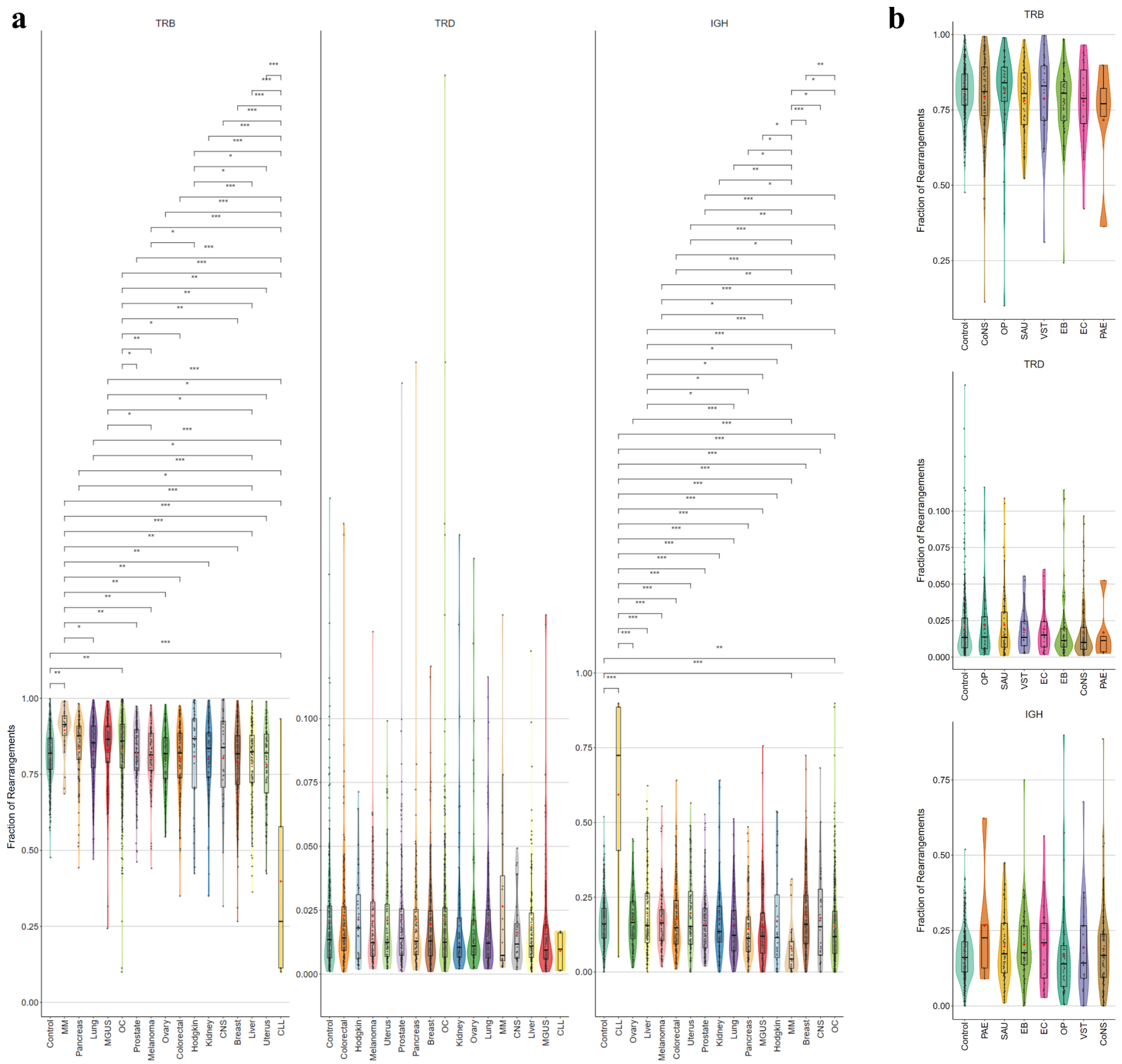


**Supplementary Figure 9**. Analysis of chain fractions based on disease status. a) Comparison of fraction of TRB, TRD, and IGH fractions in cancer subtypes and their comparison to control and each other. b) Comparison of fraction of TRB, TRD, and IGH fractions in infection subtypes and their comparison to control and each other. Statistical test was performed using a robust linear regression model (with covariate adjustment) and p values are FDR or Benjamini-Hochberg adjusted. * represents p value<0.05; ** p<0.01; *** p<0.001.

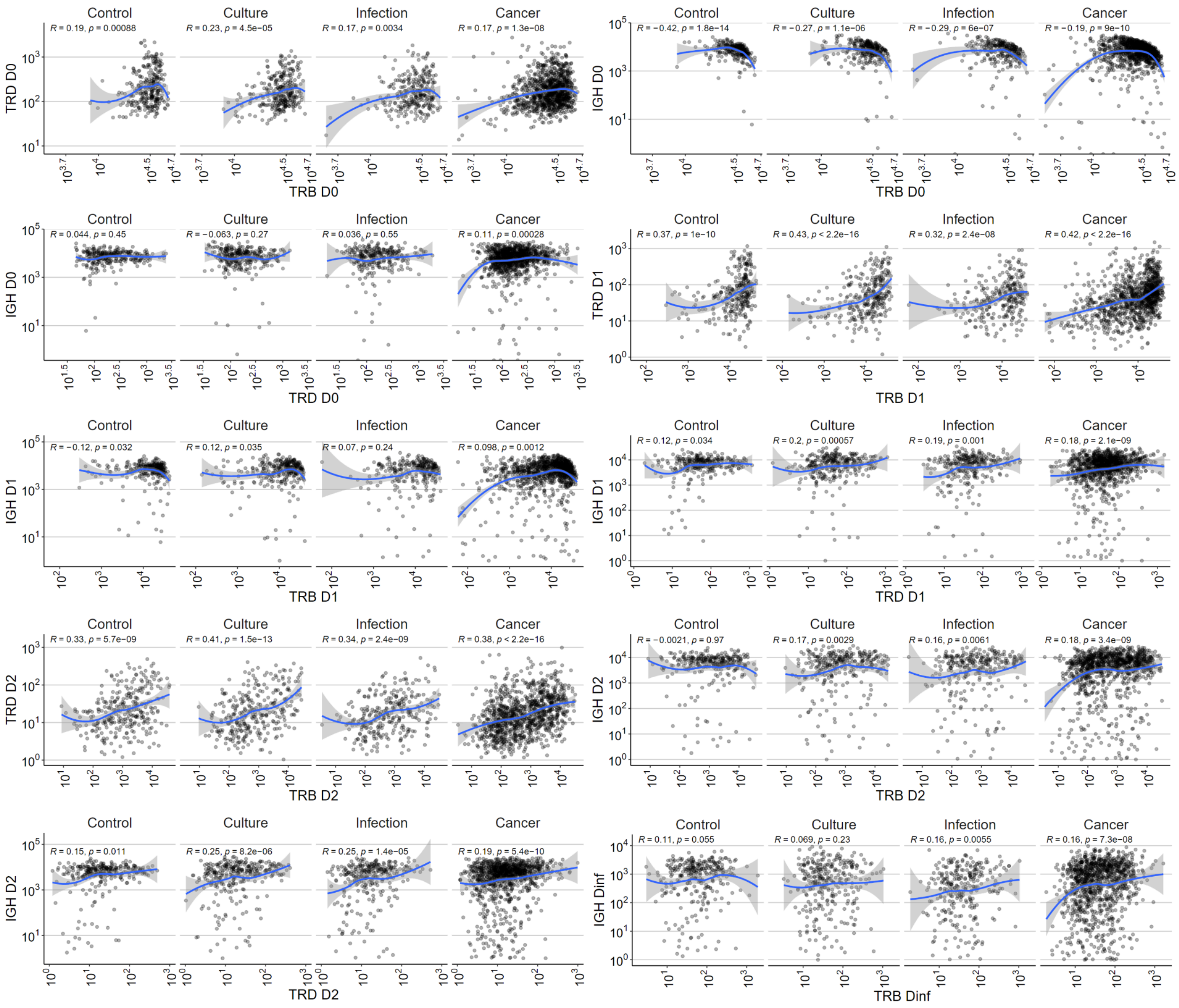


**Supplementary Figure 10**. Spearman correlation of immune repertoire diversity between different chain types in downsampled data.

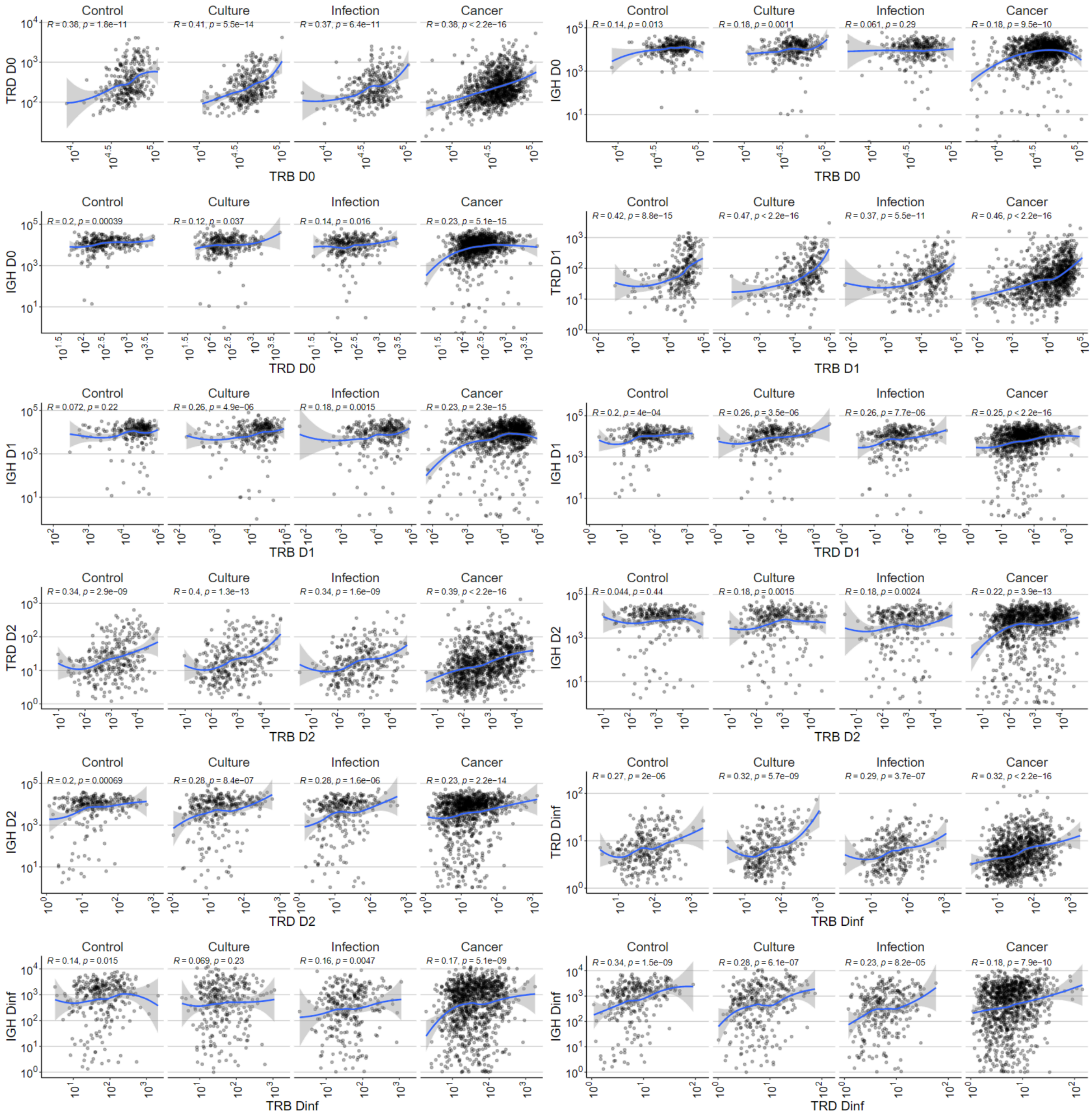


**Supplementary Figure 11**. Spearman correlation of immune repertoire diversity between different chain types in non-downsampled data.

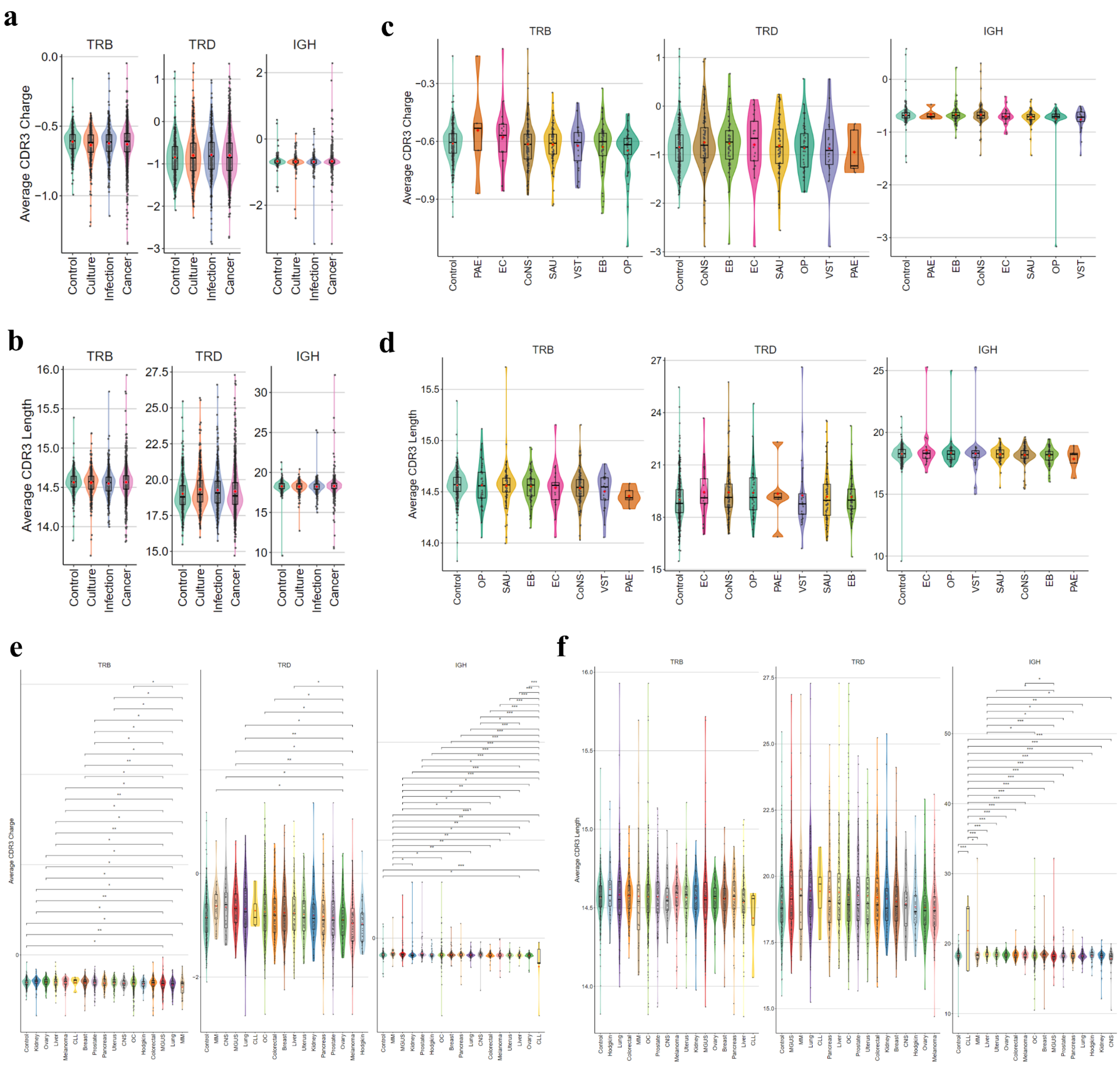


**Supplementary Figure 12**. Analysis of CDR3 length and charge with regards to disease status. a) Comparison of average CDR3 charge between conditions. b) Comparison of average CDR3 length between conditions. c) Comparison of average CDR3 charge for infection subtypes and their comparisons with control and each other. d) Comparison of average CDR3 length for infection subtypes and their comparisons with control and each other. e) Comparison of average CDR3 charge for cancer subtypes and their comparisons with control and each other. f) Comparison of average CDR3 length for cancer subtypes and their comparisons with control and each other. Statistical test was performed using a robust linear regression model (with covariate adjustment) and p values are FDR or Benjamini-Hochberg adjusted. * represents p value<0.05; ** p<0.01; *** p<0.001.

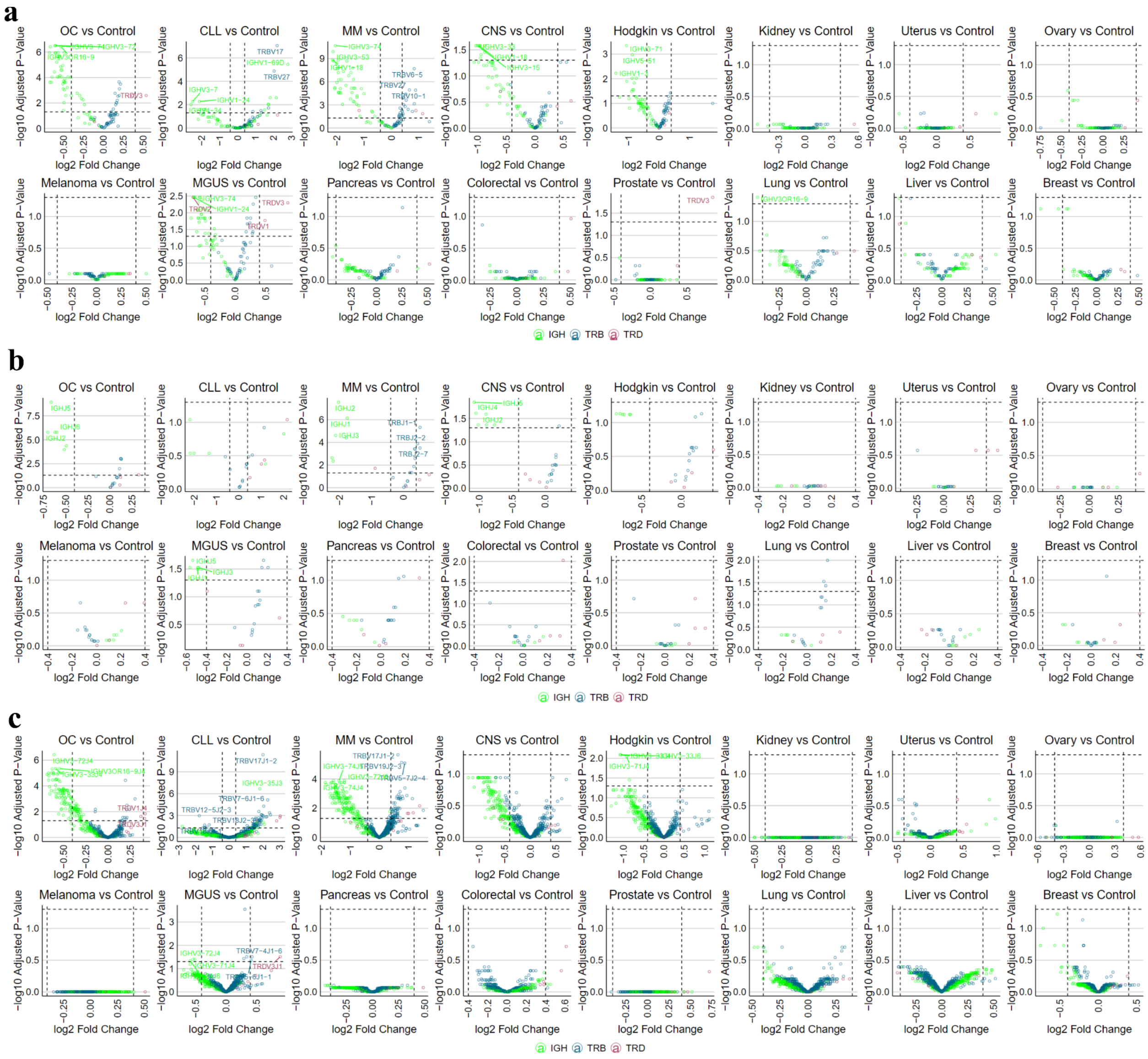


**Supplementary Figure 13**. Differential gene usage results of V (a), J (b), and VJ pair (c) Gene usage analysis for cancer subtypes and their comparison to control. Gene usage analysis was performed using limma-voom (with covariate adjustment) and p values are Benjamini-Hochberg adjusted. For the sake of visualization in volcano plots, only top 3 up- or down-used genes are labeled.

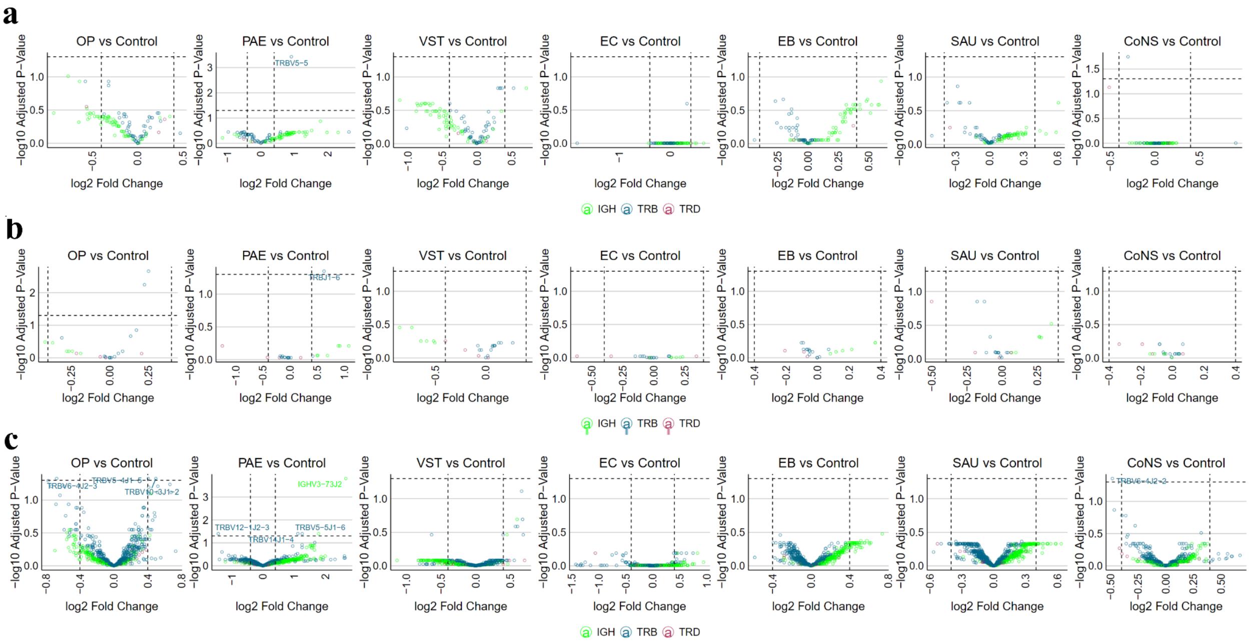


**Supplementary Figure 14**. Differential gene usage results of V (a), J (b), and VJ pair (c) Gene usage analysis for infection subtypes and their comparison to control. Gene usage analysis was performed using limma-voom (with covariate adjustment) and p values are Benjamini-Hochberg adjusted. For the sake of visualization in volcano plots, only top 3 up- or down-used genes are labeled.

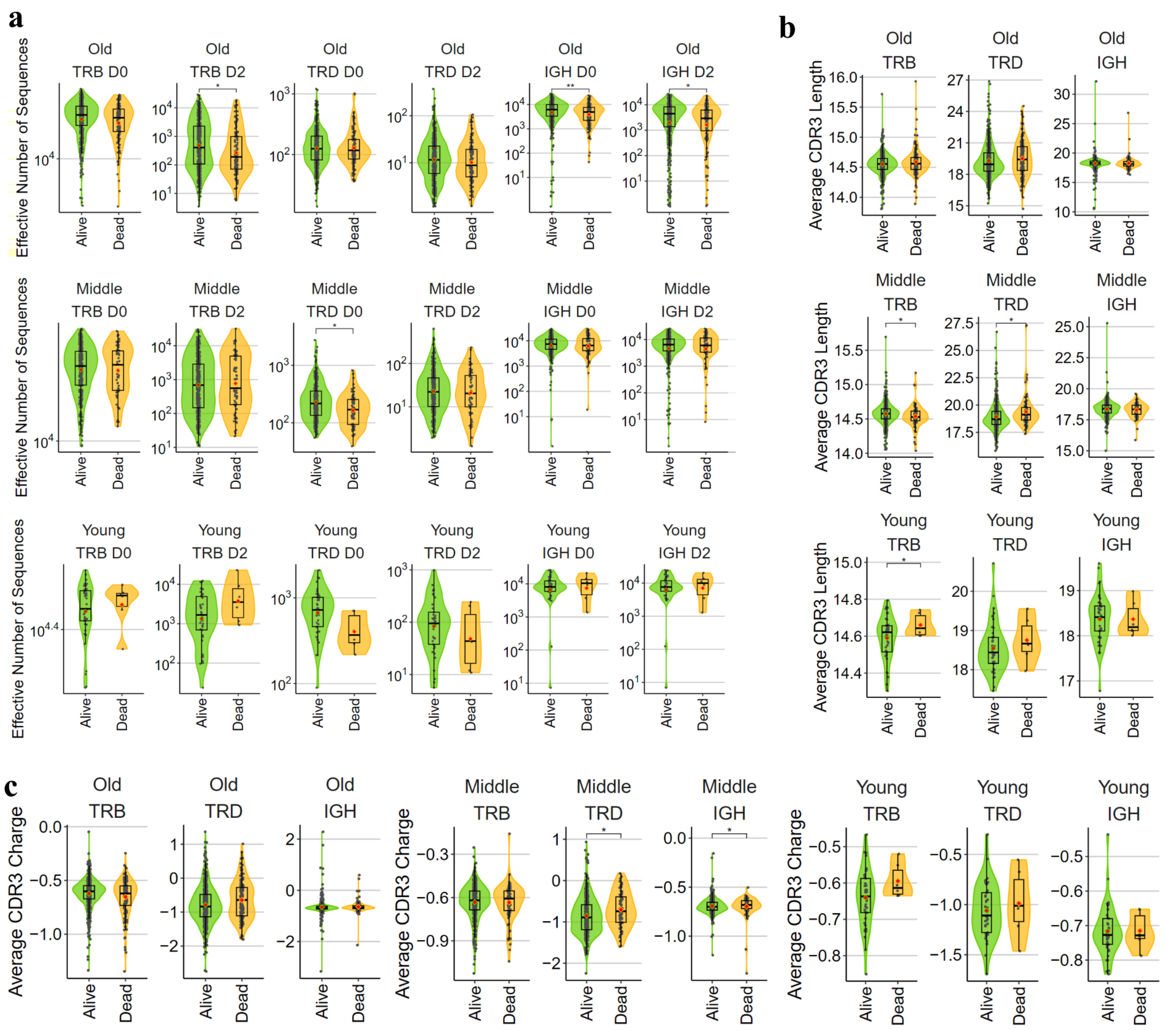

**Supplementary Figure 15**. Analysis of immune repertoire diversity with regards to survival in cancer participants. a) Comparison of $D^q$ numbers for alive and dead participants at orders 0 and 2 in three age groups. b) Comparison of CDR3 length for alive and dead participants in three age groups. c) Comparison of CDR3 charge for alive and dead participants in three age groups. Statistical test was performed using a robust linear regression model (with covariate adjustment) and p values are FDR or Benjamini-Hochberg adjusted. * represents p value<0.05; ** p<0.01; *** p<0.001. Young (<40 years old). Middle-age (40-65 years old). Old (>65 years old).

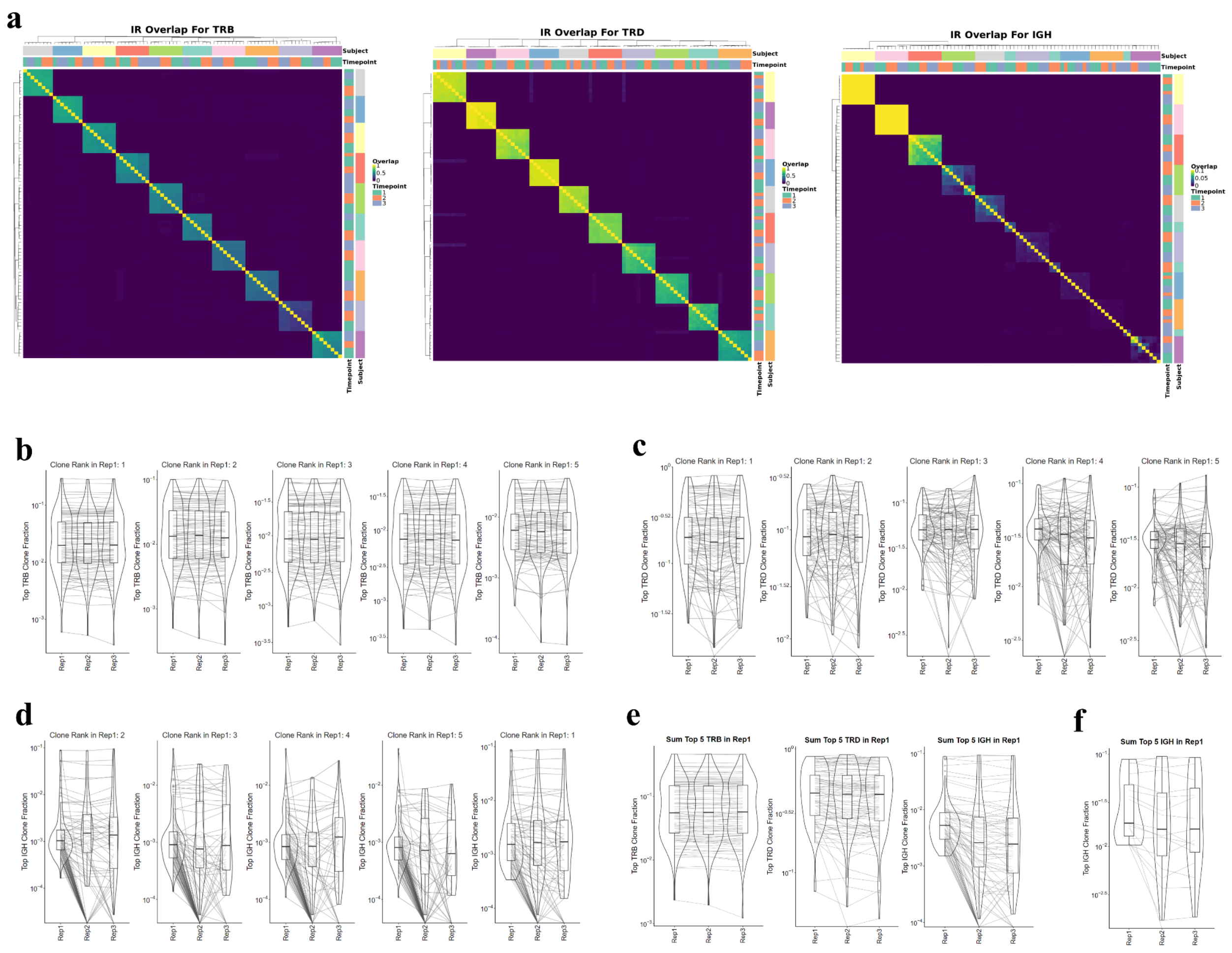


**Supplementary Figure 16**. a) Clustering of pairwise overlap of exact clonotype match for 10 randomly selected multi-timepoint participants for each chain type. b-d) Tracking clone frequency of the top 5 clonotypes across replicates of 100 randomly selected 3-replicate samples for TRB (b), TRD (c), and IGH (d). e) Tracking summed clone frequency of the top 5 clonotypes across replicates of 100 randomly selected 3-replicate samples for each chain type. f) Tracking summed clone frequency of the top 5 clonotypes if the sum is >0.01 for IGH.
